# Importin-β1 functions as a chromatin sensor to position the contractile ring for cytokinesis

**DOI:** 10.1101/2025.11.12.688041

**Authors:** C. R. Brancheriau, K. Larocque, I. Tountas, A. Perlman, A. Piekny

## Abstract

Cytokinesis, thefinal step of cell division, relies on the ingression of a carefully positioned actomyosin ring. Chromatin-associated Ran-GTP helps position this ring although the mechanism is not clear. We hypothesize that a depleted zone of Ran-GTP between the segregating chromosomes causes importins to become equatorially enriched and promote the recruitment of the scaffold protein anillin. However, the role of importins during anaphase remains unclear. Here, we tested if importins respond to the chromatin-associated Ran-GTP gradient to regulate ring assembly. Using endogenous tagging, live-cell imaging, and optogenetic perturbations, we found that importin-β1 becomes equatorially enriched and is required for cytokinesis in hypotriploid HeLa cells, but not in euploid HCT 116 cells. A predictive model of the Ran-GTP gradient and Ran-free importin-β1, validated experimentally by FLIM-FRET, identified factors that modulate this chromatin sensing pathway including chromatin-to-cell size ratio. Our findings suggest that highly aneuploid cancer cells may depend on importin-mediated anillin recruitment, representing a targetable weakness that distinguishes them from diploid cells.

**Summary blurb:** Importin enrichment at the equator promotes anillin recruitment during cytokinesis in a ploidy-dependent manner, linking Ran-GTP gradients to cell division.

**Highlights:**

- Ran-free Importin-β1 becomes enriched between segregating chromosomes in anaphase HeLa cells but not in HCT116 cells.
- Importin-binding is required for anillin localization and function in cytokinesis in HeLa cells but not in HCT 116 cells.
- Importin-β1 disruption at anaphase causes ring oscillation and cytokinesis failure in HeLa cells.
- Importin-β1 is not equatorially enriched and anillin is broader in HeLa cells with a lower chromatin-to-cell-size ratio.
- Importin-β1 is equatorially enriched and anillin is narrower in HCT 116 cells with a higher chromatin-to-cell-size ratio.

**Graphical abstract:** 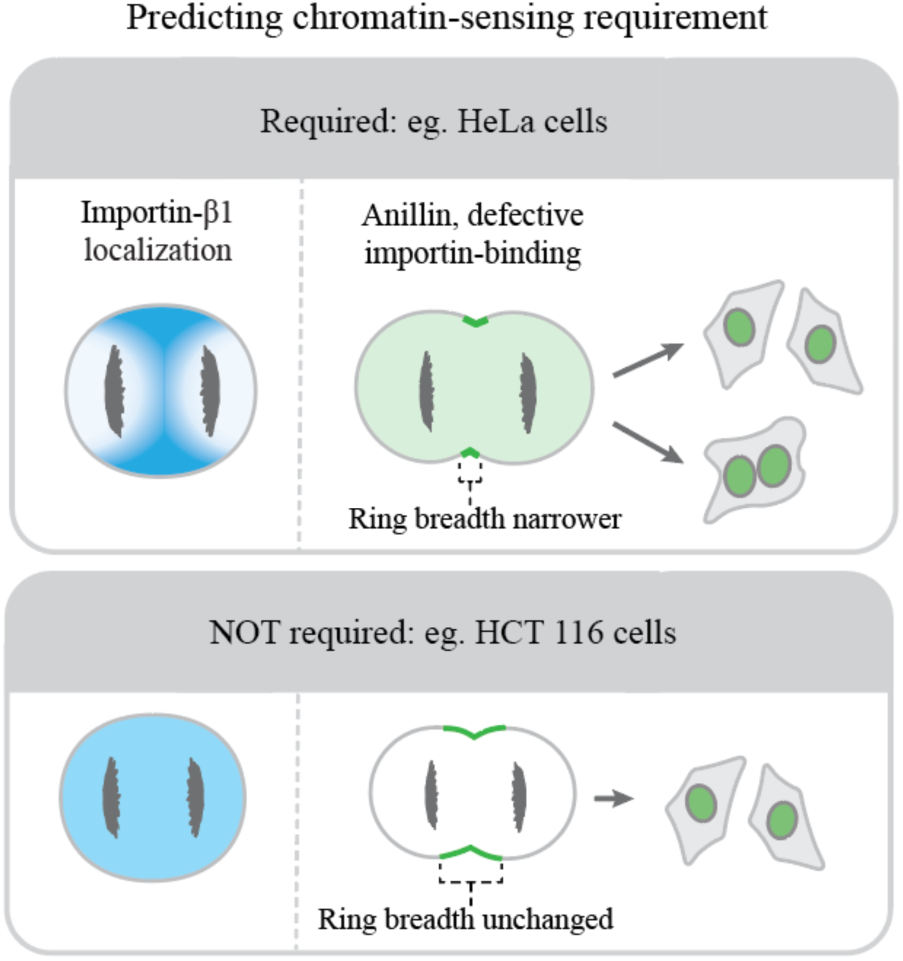

## Introduction

Cytokinesis describes the physical separation of a cell into two daughter cells. This process occurs by the ingression of a RhoA-dependent contractile ring, which is spatiotemporally regulated by multiple pathways associated with the mitotic spindle and chromatin^1,2^. Failed cytokinesis can lead to aneuploidy and altered cell fate, contributing to a range of diseases. Our group and others previously identified a chromatin-sensing pathway that coordinates contractile ring position with chromosome segregation for successful cytokinesis^3,4,5,6^.

Chromatin sensing relies on the importin-mediated binding and regulation of proteins with nuclear localization signals (NLSs) ^7,8^. The release of NLS-proteins from importins is spatially controlled by active Ran, which is enriched near chromatin since its activator RCC1 (RanGEF) is bound to histones, while RanGAP inactivates Ran in the cytoplasm^9,10,11^. Importins typically bind to NLS’s as heterodimers of importin-α and -β. While there is one predominant importin-β (β1), there are seven importin-α isoforms that are partly redundant with subfamily-specific preferences^12^. Further, importin-β1 can bind to several NLS-proteins independently of importin-α^13^. During anaphase, we predict that active Ran remains associated with the segregating chromosomes creating an equatorially depleted zone where importins are free to bind to NLS-proteins. Since several contractile proteins have NLS’s, we hypothesize that importins promote the recruitment of contractile proteins to form a well-positioned ring. In support of this hypothesis, targeting Ran-GTP to the equatorial membrane displaced ring proteins in HeLa cells, while reducing Ran-GTP levels during anaphase caused ring positioning defects in baby hamster kidney epithelial (BHK) cells^5^.

We found that the contractile ring protein anillin is a target of the Ran-importin pathway^5,6^. Anillin crosslinks actomyosin with the plasma membrane to stabilize the ring, and contains a conserved C-terminal NLS that does not control nuclear localization but is essential for anillin localization and function in HeLa cells^5,14,15,16^. Importin-β1 binds directly to anillin, and mutations that reduce importin-β1-binding impair anillin recruitment to the equatorial cortex in anaphase HeLa cells^6^. These findings suggest that importins regulate anillin to coordinate ring position with chromatin. However, the timing and breadth of anillin’s localization at the equatorial cortex differs in HCT 116 and HEK293 cells compared to HeLa cells^17^ supporting that the mechanisms regulating ring assembly change with cell type. Multiple factors could affect Ran-GTP and its ability to regulate chromatin-cortical signaling.

It is not clear which factors could affect the ability of the Ran-importin pathway to regulate the ring. Presumably ploidy could cause a change in the steepness and/or extent of the gradient of active Ran. We previously found that mild perturbation of Ran regulators or importins had different effects on ring kinetics in AB compared to P1 cells in early *C. elegans* embryos^18^. These cells have different sizes and fates. Further, increasing their ploidy from 2n to 4n caused them to increase in size by 1.5-fold, and altered both ring kinetics and reliance on Ran regulation^18^. This sublinear correlation with ploidy and cell size means that chromatin could potentially be closer to the cortex in cells with higher ploidy. Further, the levels of Ran, importins, RCC1 or RanGAP could also vary with cell type, and/or post-translational modifications could alter their localization. FRET imaging previously showed that the metaphase Ran-GTP gradient is steeper in cancer cells with higher ploidy compared to diploid HFF-1 cells^19^. A model was developed to predict the location of the active Ran gradient in metaphase HeLa cells^20^. This model revealed that the GTP/GDP ratio as well as the levels and localization of RCC1 were key parameters that determine the gradient. However, their model was not used to make predictions about active Ran in anaphase.

Importins have not been extensively studied in anaphase, yet we speculate that they respond to the Ran gradient to regulate contractile ring assembly. Here, we determined if importins play a role in cytokinesis. We chose to focus on importin-β1 given the possible redundancy among importin-α proteins. Using endogenous tagging and optogenetic perturbations, we found that importin-β1 is equatorially enriched and required for cytokinesis in hypotriploid HeLa cells, but not in euploid HCT 116 cells. These findings were supported by FLIM-FRET analysis and a predictive model of the Ran-GTP gradient during anaphase in both cell types. We then used the model to identify factors that could modulate the Ran-importin pathway’s role in cytokinesis and validated these predictions experimentally. We found that the pathway is required in cells with a higher chromatin-to-cell size ratio. Since many cancer cells have high aneuploidy with a net gain in chromosomes, their potential dependence on chromatin sensing for ring assembly may represent a therapeutic vulnerability that distinguishes them from euploid cells.

## Results

### Importin-β1 is transiently equatorially enriched in anaphase HeLa cells but not in HCT 116 cells

Importins have not been extensively studied during anaphase, but previous studies by our group suggests that they control anillin localization for cytokinesis in HeLa cells (Fig. 1A) ^5,6^. To visualize where importins localize during mitotic exit, we endogenously tagged importin-β1 with mNeonGreen using CRISPR/Cas9 in HeLa cells (origin: cervical adenocarcinoma from a Black female), and HCT 116 cells (origin: colorectal cancer from a Caucasian male; Fig. 1B and Supp. Fig. S1). As shown in Figure 1C, importin-β1 is cytosolic, but we observed a transient enrichment of importin-β1 near the equatorial cortex in anaphase HeLa cells, but not in HCT 116 cells, where the localization was more uniform. These observations were quantified using linescans drawn just under the cortex from one pole to the other during anaphase to reveal differences in fluorescence intensity in a population of cells (Fig. 1D). We also used linescans to measure fluorescence over time in HeLa cells, which revealed that the equatorial enrichment of fluorescence relative to the poles occurs between 2-6 minutes after anaphase onset and is uniform before and after this time (Fig. 1E). Importin-β1’s equatorial enrichment in HeLa cells occurred transiently as chromosomes segregated toward their respective poles (Fig. 1F). Measurements revealed that there was a positive correlation in the breadth of enriched importin-β1 and the distance between the chromosomes (Fig. 1G). These data show that importins become transiently equatorially enriched in anaphase HeLa cells as chromosomes segregate, but not in HCT 116 cells. Notably, the timing of enrichment correlates with when anillin is first visibly enriched at the equatorial cortex in HeLa cells (Supp. Fig. S2)^5,6^.

**Figure 1.**
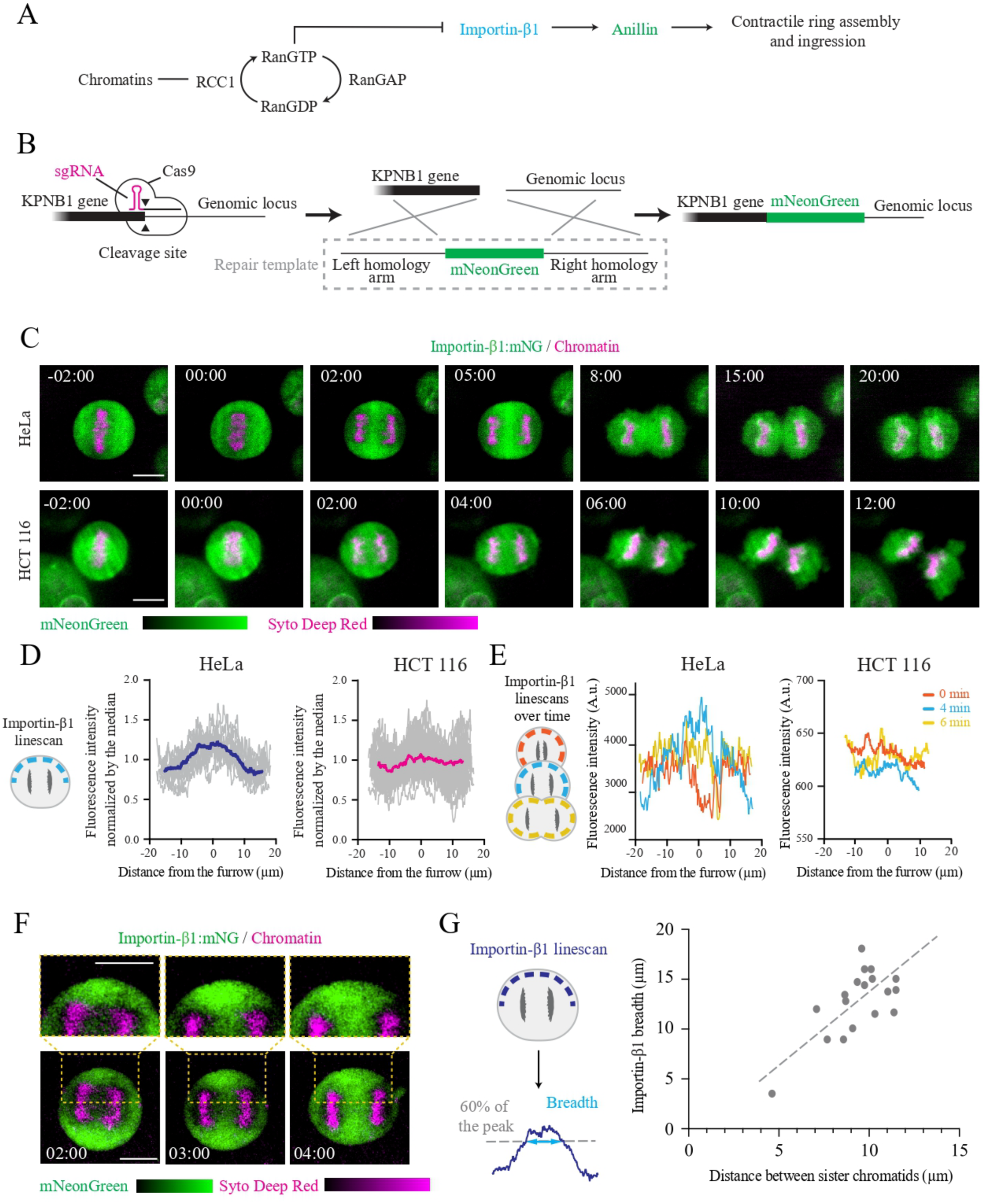
Importin-β1 is transiently equatorially enriched in anaphase HeLa cells. (A) A pathway shows how chromatin sensing via Ran-GTP could control ring assembly by the importin-mediated regulation of anillin. (B) A schematic shows how genome editing was performed in HeLa and HCT 116 cells to insert mNeonGreen at the C terminus of importin-β1. (C) Timelapse images show the localization of endogenous importin-β1 (green) co-stained for chromatin (SYTO Deep Red; magenta) in HeLa cells (top) or HCT 116 cells (bottom). The colour scales show levels (dark indicates lower fluorescence intensity compared to bright colours). Time is in minutes after anaphase onset, and the scale bar is 10 µm. (D) A cartoon cell shows how importin-β1 was measured in anaphase cells. A thick line (2 µm) was drawn subcortically from one pole to the other and the intensity along this line was plotted. Graphs show linescans of importin-β1 fluorescence in HeLa (blue) or HCT 116 (pink) cells (n=20 for each) normalized by their median over the length of the cell (shown as distance from the furrow). (E) Cartoon cells show how linescans were drawn at anaphase onset (0 min, cyan), late anaphase (4 min, orange) and early telophase (6 min, yellow). A graph shows the fluorescence intensity over distance from the furrow for a HeLa (left) and HCT 116 cell (right). (F) Timelapse images show endogenous importin-β1 (green) in a HeLa cell (bottom) from anaphase onset co-stained for chromatin (SYTO Deep Red; magenta). The images above show a zoomed-in view of the yellow boxes. Time is in minutes after anaphase onset, and the scale bar is 10 µm. (G) A graph shows how the distance between segregated sister chromatids (µm) correlates with the breadth of equatorially enriched importin-β1 (µm). The dashed line shows linear regression (r^2^= 0.43; n=20 cells).

### Importin-β1 equatorial enrichment in HeLa cells is dependent on anillin

The timing and breadth of importin-β1’s equatorial enrichment appears to correlate with anillin^5,6^. To test this, we imaged HeLa cells with endogenous importin-β1:mNG co-expressing mScarlet:anillin (Fig. 2A). Indeed, the enrichment of importin-β1 correlated with when anillin became visibly enriched at the equatorial cortex. To compare their breadth, we measured the angle between the equatorial axis and the boundary of importin or anillin enrichment in anaphase cells (Fig. 2B). Although importin-β1 enrichment was broader than anillin, there was a positive correlation between both proteins (Fig. 2B). This finding suggests that the breadth of anillin increases with the breadth of importin-β1 enrichment or vice versa.

**Figure 2.**
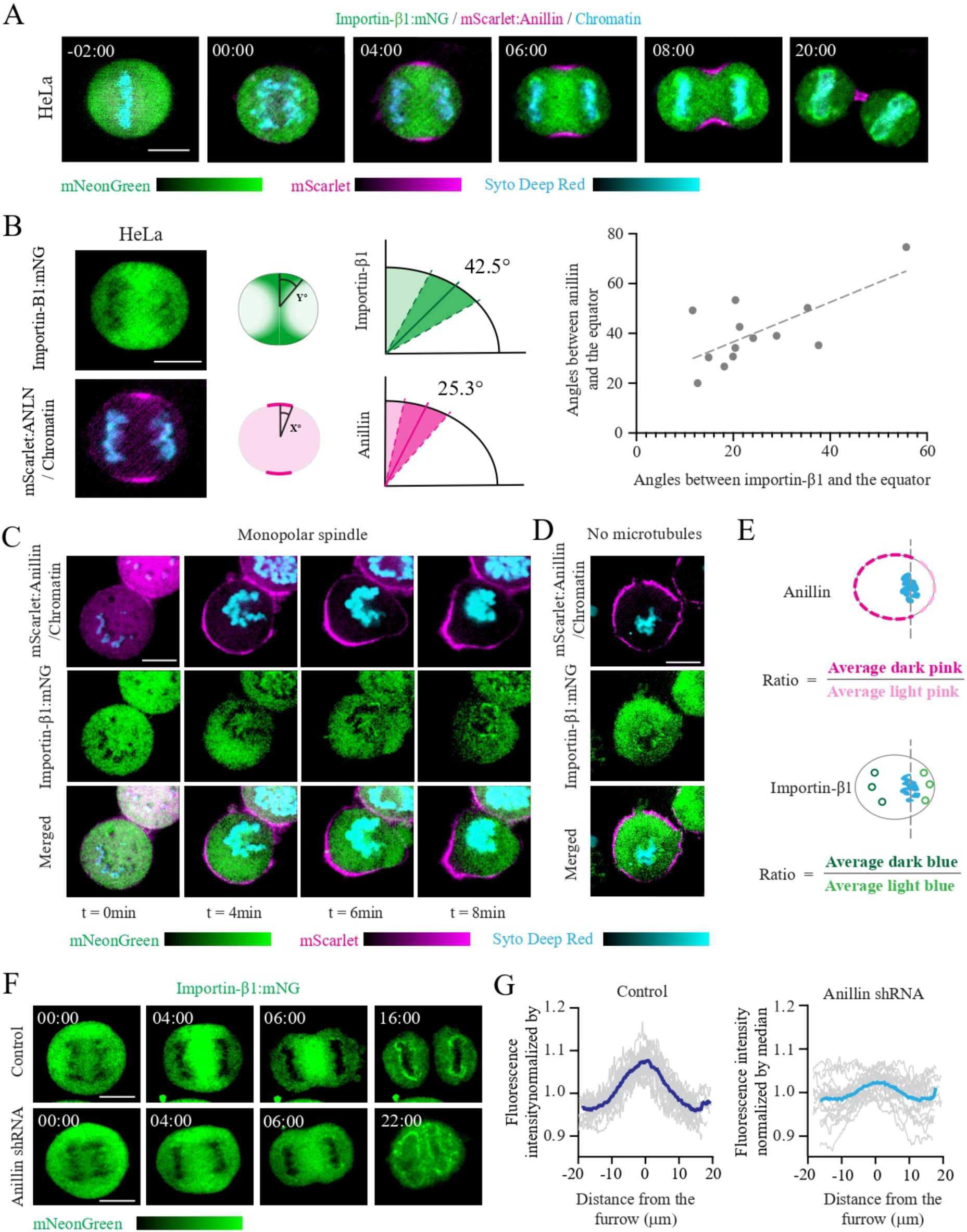
The equatorial enrichment of importin-β1 is dependent on anillin. (A) Timelapse images show endogenous mNG:importin-β1 (green) in a HeLa cell co-expressing mScarlet:anillin (magenta) and stained for chromatin (SYTO Deep Red, cyan). Time is in minutes after anaphase onset, and the scale bar is 10 µm. (B) Images show endogenous mNG:importin-β1 (green, top) in a HeLa cell co-expressing mScarlet:anillin (magenta, bottom) and stained for chromatin (SYTO Deep Rred, cyan) 4 minutes after anaphase onset. The scale bar is 10 µm. Cartoon cells show how angles between the equatorial axis and the boundary of anillin (green) and importin-β1 (pink) enrichment were measured. The origin is at the center of the cell, while the vertical axis is at the equator, and the horizontal axis is at poles. Values are indicated in the graphs to the right, where the darker colored area represents the standard deviation. A graph shows the correlation between anillin (y-axis) and importin-β1 (x-axis) enrichment. The dashed line shows linear regression (r^2^= 0.52; n= 13 cells). (C) Timelapse images show endogenous mNG:importin-β1 (green) in a HeLa cell co-expressing mScarlet:anillin (magenta) and stained for chromatin (SYTO Deep Red; cyan) induced to undergo monopolar cytokinesis. Time is in minutes after Cdk1 inhibition, and the scale bar is 10 µm. (D) Images show a single time point of endogenous mNG:importin-β1 (green) in a HeLa cell co-expressing mScarlet:anillin (magenta) and stained for chromatin (SYTO Deep Red; cyan) after microtubule depolymerization and polarization. The scale bar is 10 µm. (E) Schematics show how the symmetry of importin-β1 and anillin enrichment was measured. The average intensity of importin-β1 in regions of interest (ROI) in the cytosol away from chromatin was divided by a region near chromatin. The average intensity of a cortical linescan of anillin in a region of the cortex away from chromatin was divided by a region near chromatin. (F) Timelapse images show endogenous mNG:importin-β1 (green) in a control HeLa cell (top) or after anillin depletion by shRNAs (bottom). The colour scale shows intensity (dark indicates low fluorescence intensity compared to bright colours). Time is in minutes after anaphase onset, and the scale bar is 10 µm. (G) Graphs show linescans of importin-β1 fluorescence in control HeLa cells (dark blue) or after anillin RNAi (light blue; n=10 for each) normalized by their median over the length of the cell (shown as distance from the furrow).

We also tested if importin-β1 enrichment correlates with anillin in cells that are induced to undergo monopolar cytokinesis. We previously showed that anillin is asymmetrically enriched at the polarized cortex in cells undergoing monopolar cytokinesis, as well as in the absence of polymerized microtubules^5^. HeLa cells were treated with 2 μM of S-trityl-l-cysteine (STC) to block centrosome separation and mitotic exit was induced by inhibiting Cdk1 with 22.5 μM of Purvalanol A^21^. We observed the transient enrichment of importin-β1 near the polarized cortex away from chromatin, and where anillin became cortically enriched (Fig. 2C). We noticed that importin-β1 enrichment occurred at 4 minutes after Cdk1 inhibition, while anillin’s peak enrichment occurred at 6 minutes, with noticeable cortical ingression after 8 minutes. Importin-β1 and anillin were also enriched near the polarized cortex in HeLa cells treated with 33 nM nocodazole to depolymerize microtubules and induced to exit mitosis by Cdk1 inhibition (Fig. 2D). Next, we measured the ratio of importin-β1 and anillin enrichment on either side of chromatin in monopolar cytokinesis or after polarization of cells lacking microtubules. Both proteins showed asymmetric enrichment in relation to chromatin, which was 1.5±0.2 and 2.5±0.5 or 1.4±0.1 and 2.1±0.7 for importin-β1 and anillin, respectively (Fig. 2E). These findings support that importin-β1 becomes asymmetrically enriched away from chromatin in polarized cells, which precedes anillin’s recruitment to the nearby cortex.

Next, we tested if importin-β1’s equatorial enrichment is dependent on anillin. Importin-β1 was previously shown to localize to the metaphase spindle via its cargo, TPX2^22^. Since importin-β1 binds directly to anillin, we speculated that importin-β1’s equatorial enrichment in anaphase cells could be dependent on anillin. Indeed, depletion of anillin by expressing shRNAs in HeLa cells caused a reduction in the transient enrichment of importin-β1 (Fig. 2F and G). There is a small amount of importin-β1 enrichment that remains, which could reflect other targets that are also regulated by importin-β1 for ring assembly. This finding shows that the transient anaphase enrichment of importin-β1 is at least partly due to anillin, which is consistent with anillin being a target of importin-β1-binding.

### Importin-β1 is required for ring assembly in HeLa cells but not in HCT 116 cells

We previously found that mutating the NLS in anillin reduces importin-β1-binding and causes cytokinesis failure in HeLa cells^5^. Since importin-β1 is not equatorially enriched in HCT 116 cells, we determined if anillin’s localization and function relies on importin-β1. To do this, we depleted endogenous anillin using shRNAs and co-expressed RNAi-resistant mScarlet:anillin or mScarlet:anillin-NLS-mutant (Fig. 3A). First, we determined if importin-binding to anillin is required for cytokinesis in HCT 116 cells. Cytokinesis failure was evaluated by counting the proportion of binucleate cells in the rescued control or NLS mutant populations (Fig. 3B). While 37±7% of HeLa cells rescued with the NLS mutant failed cytokinesis compared to 6±2% for the non-mutant control or non-treated cells (4±0.5%), there was no significant change in cytokinesis failure for the NLS mutant in HCT 116 cells compared to controls (5±0.75% vs. 4±2.5 or 2.5±1% for non-treated cells). As shown in Figure 3C, anillin was not cortically localized in HeLa cells until mid-late anaphase, when it became enriched at the equatorial cortex. The NLS mutant was recruited later in anaphase and localized to a narrower region at the equatorial cortex, similar to what was previously reported^5^. In HCT 116 cells, anillin localized to the cortex prior to mitotic exit and was more broadly localized at the equatorial cortex (Fig. 3C). NLS mutant anillin was no longer cortical during metaphase and early anaphase, but by mid-anaphase was equatorially enriched with a breadth similar to control cells (Fig. 3C). We quantified these observations by measuring the cortical to cytosol ratio of anillin in metaphase HCT 116 cells and found that while this ratio was above one for control cells, the ratio was close to 1 in cells rescued with NLS mutant anillin (Fig. 3D). We also used linescans to measure the breadth to length ratio of anillin in anaphase HeLa and HCT 116 cells and found that while the ratio was reduced in HeLa cells as expected, the average breadth was similar, although more variable, in HCT 116 cells (Fig. 3E). Importantly, the localization and breadth of endogenous anillin and exogenous RNAi-resistant mScarlet:anillin co-expressed with shRNAs in HeLa and HCT 116 cells were similar suggesting that changes in localization are not caused by over-expression (Supp. Fig. S2 and Fig. 3). Overall, although importin-β1-binding is required for anillin’s cortical localization in metaphase HCT 116 cells, it is not required for regulating anillin’s localization at the equatorial cortex or for its function in cytokinesis. Thus, other pathways likely regulate anillin’s localization and function for cytokinesis in HCT 116 cells.

**Figure 3.**
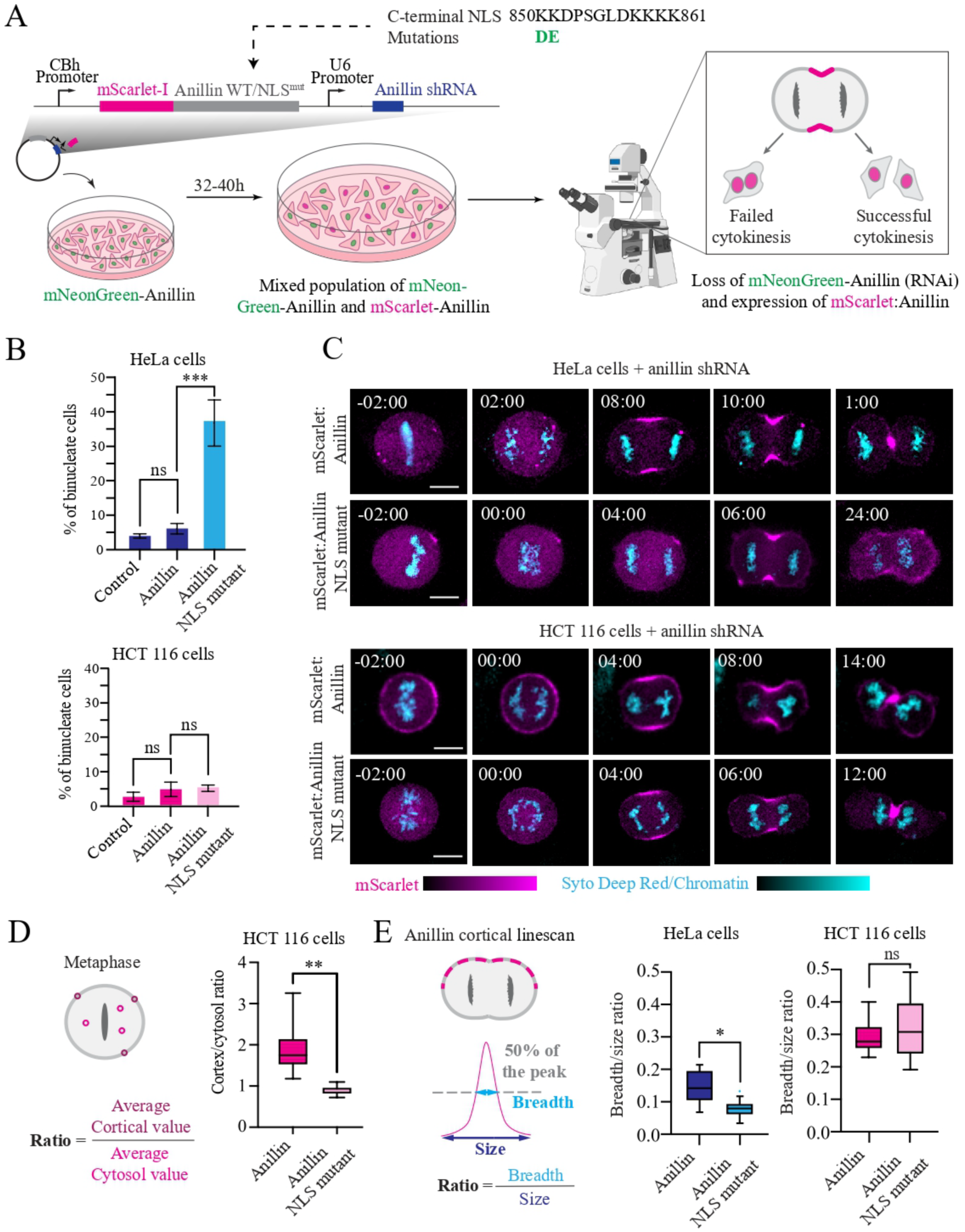
Importin-binding is required for anillin’s equatorial localization and function in HeLa cells but not in HCT 116 cells. (A) A schematic shows the workflow for the rescue experiments. Endogenous anillin was depleted by co-expressing shRNAs and RNAi-resistant anillin or the NLS mutant fused to mScarlet. Cells were imaged 32-40 hours after transfection. The point mutations in the C terminus NLS are indicated (for the anillin isoform that is 1087aa in length). (B) Bar graphs show the percentage of cytokinesis failure (binucleate cells) in non-treated HeLa (dark blue) or HCT 116 cells (dark pink) and after rescue with mScarlet:anillin (dark blue or dark pink) or mScarlet:anillin (NLS mutant; light blue or light pink; n=100 cells per replicate; N = 3). Asterisks (***) indicate significance (p ≤ 0.0001) while ns refers to non-significance. (C) Timelapse images show HeLa (top) or HCT 116 cells (bottom) rescued with mScarlet:anillin or mScarlet:anillin (NLS mutant; magenta) stained for chromatin (SYTO Deep Red; cyan). Time is in minutes from anaphase onset, and the scale bar is 10 µm. (D) A cartoon cell shows how the ratio of cortical to cytosolic anillin fluorescence was measured using three regions of interest (ROI) at the cortex and in the cytosol and dividing the averages. A bar graph shows the cortical to cytosolic ratios in metaphase HCT 116 cells (n = 15) rescued with mScarlet:anillin (dark pink) or mScarlet:anillin (NLS mutant; light pink). Asterisks (**) indicate significance p ≤ 0.001. (E) A cartoon cell shows how the breadth to length ratio was measured 6 minutes after anaphase onset. A line was drawn along the cortex from one pole to the other to plot fluorescence intensity along the line. The breadth of the anillin peak was measured as the number of pixels >50% of the maximum value, which was then divided by the total number of pixels. The bar graphs show the breadth to length ratio of mScarlet:anillin (dark blue) and mScarlet:anillin (NLS mutant; light blue) in anaphase HeLa cells (n=10) and in HCT 116 cells (n=15; dark pink and light pink, respectively). Asterisks (*) indicate significance (* p ≤ 0.005) while ns refers to non-significance.

Next, we wanted to determine if importin-β1 is required for ring assembly in HeLa cells. Since importin-β1 has pleiotropic functions^8,22^, we designed an optogenetic tool to disrupt its function with high temporal precision. The optogenetic tool expresses importin-β1 fused to the fluorescent protein mCherry and CRY2clust (Fig. 4A). CRY2clust is a modified version of CRY2 with enhanced homo-oligomerization^24^. Clustering of CRY2clust was previously shown to occur rapidly in response to 400-500 nm of light^24,25,26^. HeLa cells with endogenous importin-β1:mNG were transfected with importin-β1 siRNAs to deplete the endogenous protein while co-expressing RNAi-resistant importin-β1:mCherry:CRY2clust (Fig. 4). Since exposure to 488 nm can activate CRY2clust and excite mNeonGreen, the tool was activated while imaging for the depletion of endogenous importin-β1 at anaphase onset (Fig. 4A). Cells with ≥70% depletion of endogenous importin were used for analysis. Since high over-expression of importin-β1 can affect anillin localization^5^, cells were selected with mid to low expression of importin-β1:mCherry:CRY2clust. When not activated, importin-β1:mCherry:CRY2clust localized similar to endogenous importin-β1 (Fig. 4B). When the tool was activated at anaphase onset by exposure to 488 nm of light, importin-β1 clustered with oscillation of the contractile ring, and 2 out of 5 cells failed cytokinesis (Fig. 4B). These results suggest that importin-β1 is required to stabilize the ring for cytokinesis.

**Figure 4.**
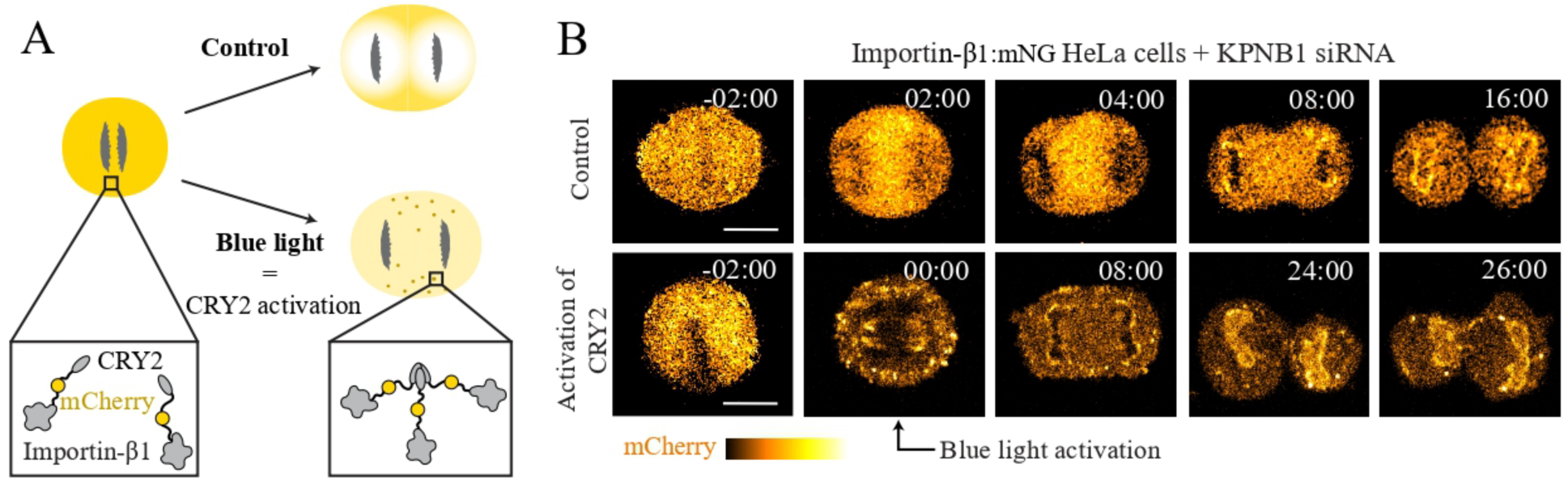
Importin-β1 stabilizes the contractile ring. (A) A cartoon schematic shows the design and regulation of importin-β1:mCherry:CRY2clust. Before activation, importin-β1:mCherry:CRY2clust is equatorially enriched, and clusters after exposure to 488 nm of light at anaphase onset due to oligomerization. (B) Timelapse images show the localization of importin-β1:mCherry:CRY2clust in a HeLa cell with endogenous importin-β1:mNG treated with importin-β1 RNAi cell during cytokinesis (orange hot LUTs: black indicates low fluorescence and yellow indicates high fluorescence). The top panel shows a control cell and the bottom panel show a cell that was activated by exposure to 488 nm of light at anaphase onset. Time is in minutes from anaphase onset, and the scale bar is 10 µm.

### Ran-GTP free importin-β is equatorially enriched in HeLa cells but not in HCT 116 cells

Our data shows that importin-β1 is equatorially enriched and controls anillin’s localization for ring assembly in HeLa cells, but not in HCT 116 cells. Importins should be free to bind to NLS-proteins where active Ran is low. To reveal where Ran-free importin-β1 is localized in HeLa and HCT 116 cells, FLIM (Fluorescence Lifetime Imaging Microscopy) was performed using Rango-3, a FRET probe designed by the Kálab laboratory^27,28,29,30^. Rango-3 has EGFP as the donor and dsREACh (non-fluorescent) as the acceptor, and the two proteins are separated by the IBB domain from snurportin, which binds to importin-β1^27^. The lifetime of EGFP decreases when FRET occurs, as photons are transferred to dsREACh. When importin-β1 is bound to the IBB domain, FRET cannot occur between EGFP and dsREACh. Consequently, regions with low lifetime indicate active FRET and *low levels* of Ran-free importin-β1, whereas regions with higher lifetime reflect the absence of FRET and *greater levels* of Ran-free importin-β1 (Fig. 5A). The lifetime of Rango-3 was assessed using the phasor plot generated by fast-FLIM and highlighting a specific region of the plot to capture the lifetime variation (Supp. Fig. S3). As shown in Supplemental Figure S4 and Supplemental Table S1, we observed high variation in the EGFP lifetime measurements among cells due to differences in the expression levels of Rango-3. Specifically, the lifetime became more uniform and decreased with higher expression of the Rango-3 probe, likely due to saturation of importin-β1-binding. Given the challenges in imaging cells with low Rango-3 expression, cells where the Rango-3 signal was suMicient for acquisition and image analysis could have been close to saturation, underestimating the levels of Ran-free importin-β1. First, we performed FLIM of Rango-3 in metaphase HeLa and HCT 116 cells and measured the distribution of lifetime changes along a radius from chromatin to the cortex (Fig. 5B and C). As expected, there was a comparable change in lifetime from the chromatin toward the cortex in both cell types indicating a switch from low to high Ran-free importin-β near the cortex (Fig. 5B and C). However, in HCT 116 cells, the overall lifetime was shorter (1.23-1.33ns) suggesting lower concentrations of Ran-free importin-β1 compared to HeLa cells (Fig. 9B and C). Next, we imaged Rango-3 in anaphase cells and measured the changes in lifetime along a thick line drawn subcortically from one pole to the other (Fig. 5B and C). In HeLa cells we saw longer lifetimes in the equatorial region (1.43ns) compared to the poles (1.35ns), while the lifetimes were shorter in HCT 116 cells and more uniformly distribution (1.23-1.3ns). Thus, there are overall lower levels of Ran-free importin-β1 that are more uniformly distributed in HCT 116 cells compared to Hela cells, which have a high pool of Ran-free importin-β1 in the equatorial region.

**Figure 5.**
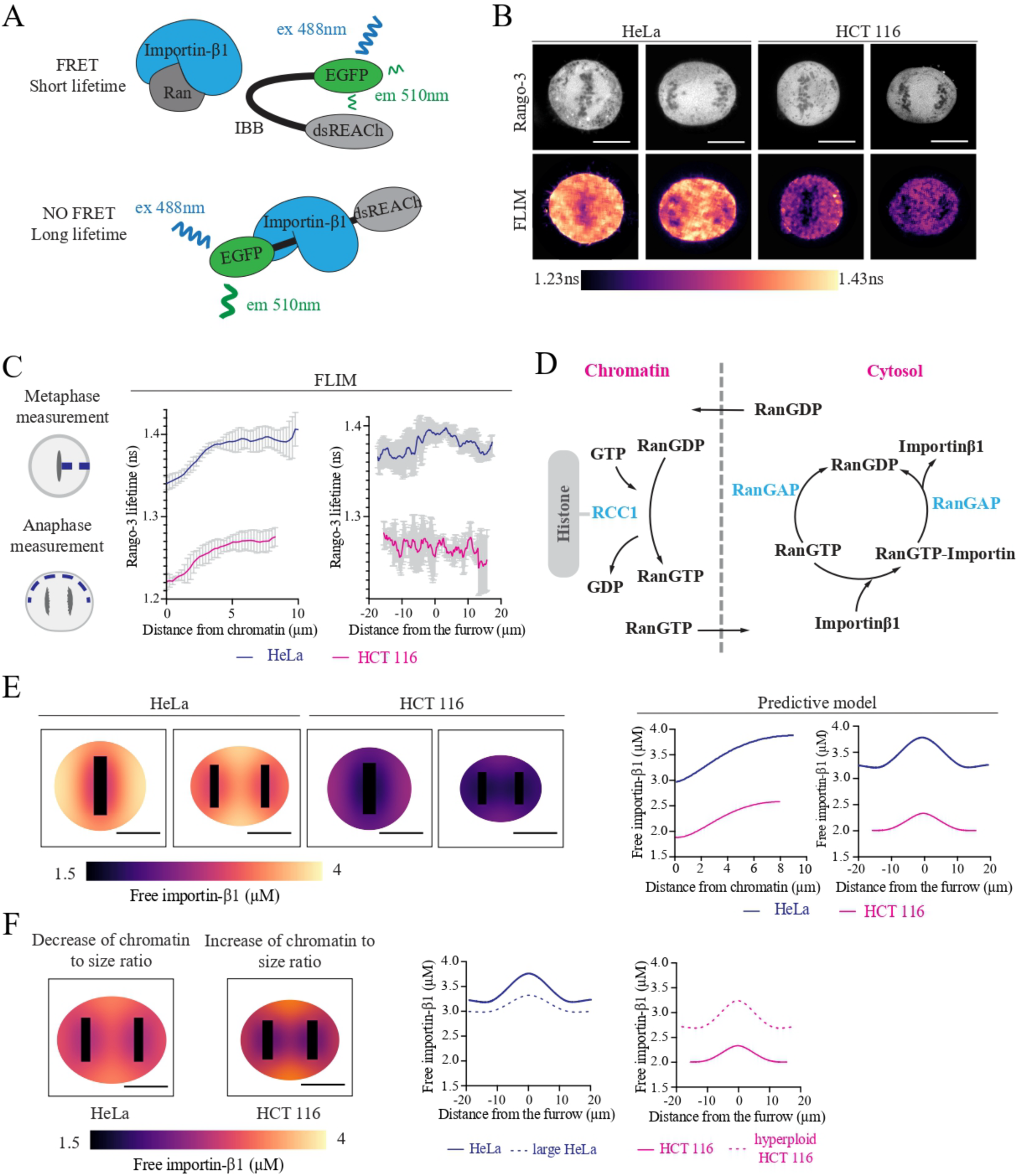
Ran-free importin-β1 is equatorially enriched in anaphase HeLa cells but not in HCT 116 cells. (A) A schematic shows the Rango-3 probe used to measure Ran-free importin-β1 in cells. EGFP is fused to the importin-β binding domain (IBB) and dsREACH, a non-fluorescent acceptor. Binding of importin-β to the IBB prevents FRET via the transfer of photons from EGFP to dsREACH, resulting in higher EGFP lifetime compared to when importin-β is not bound. (B) Images show Rango-3 fluorescence (grey, top panel) and FLIM (ns; magma LUTs: black indicates short lifetime; pale yellow indicates long lifetime) in HeLa and HCT 116 cells at metaphase and anaphase. The scale bar is 10 µm. (C) Cartoons on the left show where lines were drawn in metaphase and anaphase cells to measure lifetime from FLIM data or concentration from the models. The graphs show lifetime (ns) over distance (µm) from chromatin (metaphase) or the furrow (anaphase) for HeLa (blue) and HCT 116 cells (pink). (D) The reactions used for the computational model are shown. The model solves 8 partial differential equations in a metaphase or anaphase cell. Chromatin and cytosol are separated by a fictive limit which Ran-GTP and Ran-GDP can cross freely. RCC1 and relevant complexes are fixed in the chromatin subdomain. Other species have their own diffusion coefficients and are in the cytosol. (E) Images show the predicted levels of Ran-free importin-β1 (µM; magma LUTs: black indicate low concentrations; pale yellow indicates high concentrations) in metaphase and anaphase HeLa and HCT 116 cells. The graphs on the right show Ran-free importin-β concentration (µM) over distance (µm) from chromatin (metaphase) or the furrow (anaphase) for HeLa (blue) and HCT 116 cells (pink). (F) Images show the predicted levels of Ran-free importin-β1 (µM; magma LUTs: black indicate low concentrations; pale yellow indicates high concentrations) in anaphase HeLa and HCT 116 cells after increasing cell volume and increasing ploidy, respectively. The graphs show the changes in concentrations in Ran-free importin-β1 (μM) over distance from the furrow in anaphase HeLa (blue lines) and HCT 116 cells (pink cells) after increasing cell volume and increasing ploidy, respectively, compared to control cells.

Next, we generated a computational model to predict the Ran-free importin-β1 gradient in metaphase and anaphase HeLa and HCT 116 cells derived from a previously published 1-D model^20^. As described in the methods, we used the partial differentiation equation toolbox in MATLAB to adapt the model to a 2-D geometry which represents a cell in metaphase or anaphase. The model works by solving a series of diffusion-reaction equations using the concentrations of Ran, RanGAP, RCC1, importin-β1 and GTP/GDP, reaction rates, kinetic constants and diffusion rates in defined zones: 1) chromatin, where RCC1-mediated exchange of GDP for GTP activates Ran, and 2) cytosol, where RanGAP hydrolyzes Ran-GTP, and Ran-GTP and importin-β1 can form complexes (Fig. 5D, Supp. Fig. S5 and Supp. Table S2 and S3). To generate a gradient, we artificially sequestered RCC1 complexes in the chromatin zone, while RanGAP and importin-β1 complexes were sequestered in the cytosolic zone, and only Ran-GTP and Ran-GDP could cross the artificial boundary (Fig. 5D and Supp. Fig. 5). To determine values for the geometrical parameters, we measured length and width of the cell and chromatin (Supp. Fig. S6 and Supp. Table S4). The biochemical parameters and protein concentrations for HeLa cells were taken from previous studies (Supp. Tables S2 and S3)^20^. We used publicly available RNA sequencing data comparing HeLa and HCT 116 cells^31^ to make an approximation of the protein concentrations in HCT 116 cells (Supp. Fig. S7 and Supp. Table S5). To refine the predictive model, we solved the system of reaction-diffusion equations for HeLa and HCT 116 cells in metaphase and anaphase cells and compared the results to the experimental data, then made minor adjustments to the protein concentrations, while respecting measured values (Supp. Fig. S8 and Supp. Table S6). Among the conditions tested, we found that the concentrations of Ran and importin-β1 had greater impacts on the Ran-free importin-β1 gradient compared to RCC1 and RanGAP (Supp. Fig. S9). Using these parameters, the model predicted trends for the localization of Ran-free importin-β1 that were similar to what was obtained by the FLIM data (Fig. 5C, E and F). There was an overall lower level of Ran-free importin-β1 in HCT 116 cells compared to HeLa cells, and there were higher levels of Ran-free importin-β1 in the equatorial region of anaphase HeLa cells compared to the poles, while the distribution was more uniform in anaphase HCT 116 cells (Fig. 5C and F). Thus, our model provides a good estimate of the measured levels of Ran-free importin-β1 in cells.

### Chromatin to size ratio aGects the equatorial enrichment of Ran-free importin-β1 and anillin

We identified protein concentrations that accurately predicts the Ran-free importin-β1 gradients in metaphase and anaphase HeLa and HCT 116 cells. Another parameter that could affect the Ran-GTP gradient is the chromatin to cytosolic ratio. The steepness of the gradient is greater in metaphase cells with higher ploidy, which could impact the mechanisms regulating spindle assembly^19^. In anaphase cells, a steeper gradient could cause Ran-free importin-β1 to become equatorially enriched and regulate ring assembly. The relationship with ploidy and cell size is sublinear, where an increase in ploidy by a factor of 2 causes a 1.5 increase in cell size^32^. Thus, as ploidy increases, there is a greater chance for active Ran to reach the cortex and create differences in the equatorial enrichment of importin-β1. To test this, we used the computational model to predict how increasing the ploidy of HCT 116 cells impacts the distribution of Ran-free importin-β1 during anaphase (Supp. Table S6 and S7). The model predicted an equatorial enrichment of Ran-free importin-β1, similar to what we observe for HeLa cells (Fig. 5F). We also speculated that increasing cytoplasmic volume in HeLa cells to reduce the chromatin to cytosol ratio could have the opposite effect and reduce the equatorial enrichment of Ran-free importin-β1. Indeed, using the model to predict the impact of increasing cell volume relative to ploidy in anaphase HeLa cells resulted in a more uniform distribution of Ran-free importin-β1 (Fig. 5F).

Next, we determined if experimentally altering the chromatin to cytosolic ratio in HeLa and HCT 116 cells could affect importin-β1 and anillin localization. To increase the ploidy of HCT 116 cells, they were treated with CoCl_2_ as described in the methods (Supp. Fig. S10) ^33,34^. The volume of HeLa cells was increased using hypotonic media supplemented with FBS (Supp. Fig. S10). Spindle size scales with cell size and should not be significantly altered by our treatments^35^. However, increasing volume using hypotonicity could alter the critical concentration of free tubulin, which could impact microtubule length. Imaging microtubules in control and hypotonic HeLa cells revealed that there were no obvious differences in their spindles (Supp. Fig. S11). In hypotonic HeLa cells, importin-β1 was more uniformly distributed during anaphase, while there was an equatorial enrichment of importin-β1 in hyperploid HCT 116 cells (Fig. 6A). We measured the levels of importin-β1 relative to the furrow and indeed, there was no change in the distribution of importin-β1 in hypotonic HeLa cells, which remained similar over time (Fig. 6B and C). In hyperploid HCT 116 cells, there was a measurable increase in importin-β in the equatorial region, which appeared at 4 minutes after anaphase onset, similar to control HeLa cells (Fig. 6B and C). Changing the chromatin to cytosol ratio also had an impact on anillin localization. In hypotonic HeLa cells, anillin was cortically localized during metaphase, and became more broadly localized at the equatorial cortex during anaphase compared to control cells (Fig. 6D-H). In hyperploid HCT 116 cells, anillin was not cortically localized in metaphase, and localized to a narrower region at the equatorial cortex during anaphase compared to control cells (Fig. 6D-H). These results show that the chromatin to cytosol ratio affects the localization of importin-β1 and cortical anillin.

**Figure 6.**
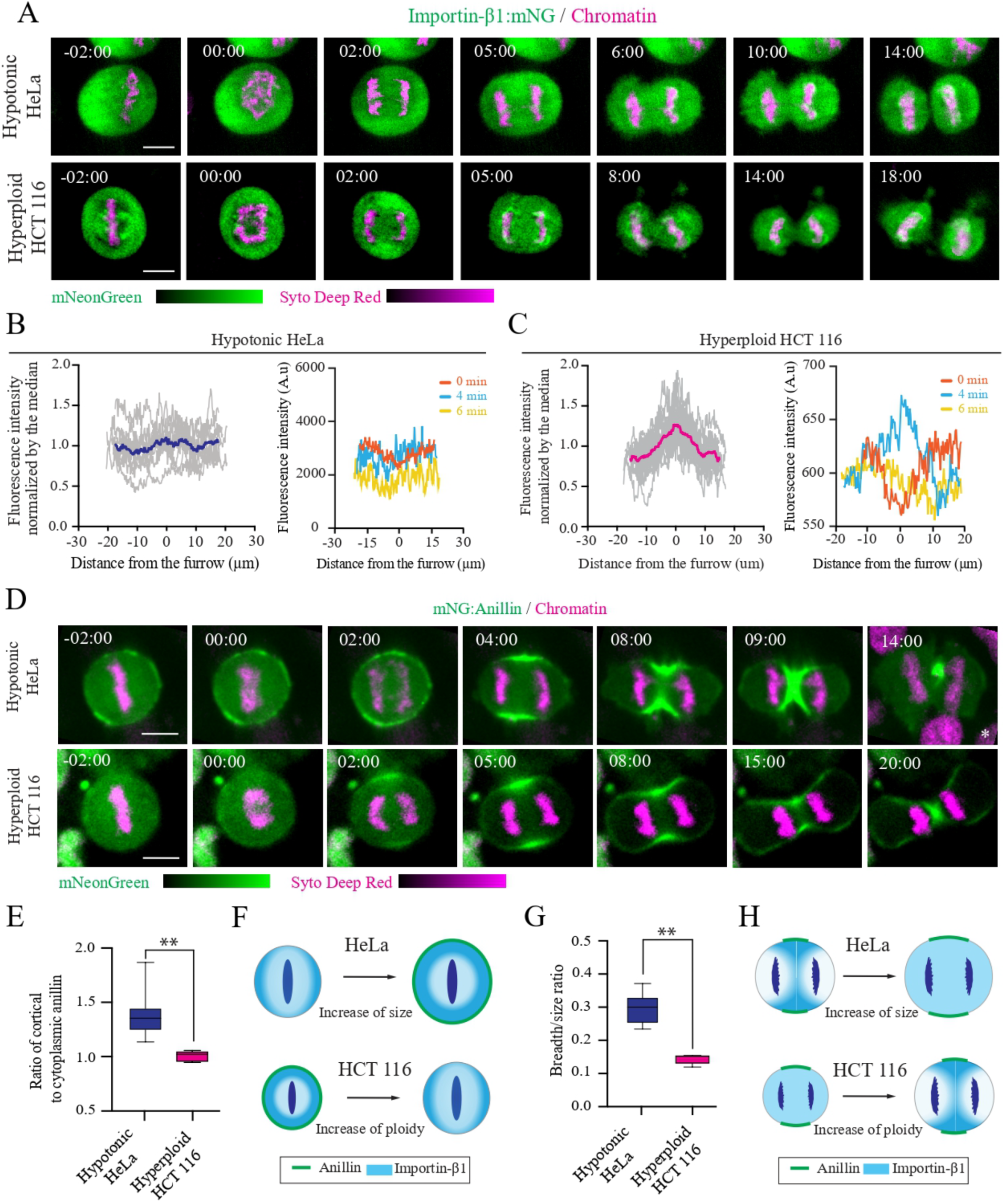
Chromatin to size ratio affects the localization of importin-β1 and anillin. (A) Timelapse images show endogenous mNG:importin-β1 (green), co-stained for DNA (SYTO Deep Red; magenta) in hypotonic HeLa cells (top) or hyperploid HCT 116 cells (bottom). Time is in minutes after anaphase onset, and the scale bar is 10 um. (B) Graphs show importin-β1 fluorescence intensity at 4 minutes after anaphase onset (left) and over time (right; 0 min in red, 4 min in blue, 6 min in yellow) in hypotonic HeLa cells (blue; n=10), and (C) in hyperploid HCT 116 cells (pink; n=10). (D) Timelapse images show endogenous mNG:Anillin (green), co-stained for DNA (SYTO Deep Red; magenta) in hypotonic HeLa cells (top) or hyperploid HCT 116 cells (bottom). Time is in minutes, and the scale bar is 10 μm. (E) Graphs show the average cortical to cytosolic ratio for anillin in metaphase hypotonic HeLa cells (blue; n=10) and hyperploid HCT 116 cells (pink; n=15; **: p ≤0.001). Bars indicate standard deviation. (F) Cartoon cells show the changes in cortical anillin in HeLa and HCT 116 cells after treatment to increase size or ploidy, respectively (anillin, green; importin-β1, blue). (G) Graphs indicate the anillin breadth to size ratio in anaphase hypotonic HeLa (blue; n=10) and hyperploid HCT 116 cells (pink; n=15; **: p ≤0.001). Bars indicate standard deviation. (H) Cartoon cells show the changes in anillin breadth in anaphase HeLa and HCT 116 cells after treatment to increase size or ploidy, respectively (anillin, green; importin-β1, blue).

## Discussion

Here we show that importin-β1 regulates anillin’s cortical recruitment during mitosis, and depending on the cell type, controls the enrichment of equatorial anillin to coordinate ring position with the segregating chromosomes. While previous data suggested that importins regulate anillin function for ring assembly in response to Ran^3,5,6,18^, importins themselves had not been extensively studied during anaphase. In HeLa cells, we show that both the total and Ran-free pools of importin-β1 become transiently equatorially enriched between the segregating chromosomes and correlates with where anillin accumulates at the overlying equatorial cortex. Importin-β1 was previously reported to have localization patterns that reflect its cargo during mitosis^22^, and we found that importin-β1’s equatorial enrichment is anillin-dependent. Disruption of importin-β1 localization during anaphase caused the ring to oscillate, similar to anillin depletion, and point mutations in the NLS of anillin that disrupt importin-binding caused cytokinesis failure in HeLa cells^5^. However, despite this requirement for importin-binding in HeLa cells, importin-β1-binding is not required for anillin localization or function in cytokinesis in HCT 116 cells. Additionally, both the total and Ran-free pools of importin-β1 are not enriched in the equatorial region of anaphase HCT 116 cells. HeLa and HCT 116 cells differ in cell fate as well as ploidy. Changing the concentrations of Ran and importin-β1 or altering the chromatin to cytosol ratio using our computational model predicted differences in the equatorial enrichment of Ran-free importin-β1. Experimentally changing the chromatin to cytosol ratio by changing ploidy or cytoplasmic volume caused both importin-β1 and anillin localization in HeLa and HCT 116 cells to change and resemble the other cell type. Thus, these parameters could be used to predict when the Ran-importin pathway plays a critical role in chromatin sensing for ring assembly during cytokinesis.

Our model was inspired by a previously published 1D model predicting the Ran gradient in metaphase cells^20^ and extending it to a 2D representation of both metaphase and anaphase cells. The Ran-free importin-β1 gradient was selected as the model output, which was confirmed using FLIM-FRET imaging of Rango-3. While our model made predictions on the levels and distribution of Ran-free importin-β1 that reflected the FLIM data, the prediction showed more enrichment in the equatorial plane of HeLa cells than what was observed. As previously mentioned, Rango-3 is likely close to saturation, which could dampen the equatorial values. In addition, our model does not include importin-α, which could cause differences in the distribution of importin-β1^12,13,36,37,38^. Further, importin-β1 localization is influenced by TPX2 in metaphase^22^, and anillin in anaphase, which were not considered in the model. The levels of anillin could define a threshold of importin-β1 that is required for regulation, and future improvements to the model could incorporate anillin binding affinities to predict this. Given that predicted Ran-free importin-β gradients in metaphase and anaphase HeLa and HCT 116 cells are similar to what was found experimentally, our findings support that chromatin to size ratio is one of the key factors that could be used to predict whether importin-β1 is required for cytokinesis. Further, since the total pool of importin-β1 localizes similar to the Ran-free pool, importin-β1 localization could be used as a proxy for predicting the role of the Ran- importin-β1 pathway in cytokinesis.

As highlighted above, our study did not test a role for importin-α in cytokinesis. The classical bipartite NLS in anillin is predicted to bind to a heterodimer of importin-α and importin-β1^37^. Our studies focused on importin-β1 because there are multiple importin-α isoforms that could bind to classical NLS’s, while importin-β1 is predicted to be the main importin-β^13,38^. Depending on the importin-α isoform, post-translational modifications that influence localization or stability could affect the levels of the heterodimer and their location, which could also impact their role in ring assembly^36^. It will be important to investigate which importin-α binds to importin-β1 to regulate cytokinesis and/or if importin-β1 functions separately.

Our model and experimental data revealed that several factors can affect the Ran-importin pathway and its ability to regulate the ring during cytokinesis. A previous study showed that chromosomal gain increases the levels of chromatin-associated RCC1 and the steepness of the Ran gradient, which affects spindle assembly in cultured human cells^19^. This steeper gradient is predicted to remain with the segregated chromosomes in anaphase, creating a greater differential enrichment of equatorial importin-β1 compared to cells with less steep gradients. Interestingly, changing RCC1 alone in the model did not have as great an impact as changing Ran or importin-β1 levels, indicating that changes in the concentration of Ran components likely also contribute to the steepness of the gradient.

Our findings support that the requirement for the Ran-importins pathway in regulating the ring for cytokinesis varies with cell type. This is consistent with our previous findings studying this pathway in *C. elegans* embryos. We found that increasing ploidy or changing the levels of active Ran by depletion of RCC1, RanGAP, importin-α and/or -β caused changes in ring kinetics, but there were differences depending on the cell type^18^. AB cells are larger and give rise to multiple tissues of the adult worm, while P1 cells are smaller and give rise to the germline. Interestingly, we found that anillin was not a target of the Ran pathway in P1 cells, suggesting that it has other targets that affect ring assembly^18^. Thus, differences in size and fate could support different requirements for the cytokinetic machinery including regulation by Ran-importins in distinct cell types. Determining which cell types rely on this pathway, and how threshold levels of anillin or other effectors influence pathway engagement will be an important direction for future studies. Given that many progressive cancers have aneuploidy with a net gain in chromosome content, targeting the chromatin-sensing pathway or its mediators such as importin-β1 may provide new strategies to restrict their proliferation.

In summary, we uncover a central role for importin-β1 as a readout for chromosome position and Ran activity to coordinate ring position via anillin, and we define the chromatin-to-size ratio as a key determinant of when cells require the chromatin-sensing pathway for successful cytokinesis.

## STAR★Methods

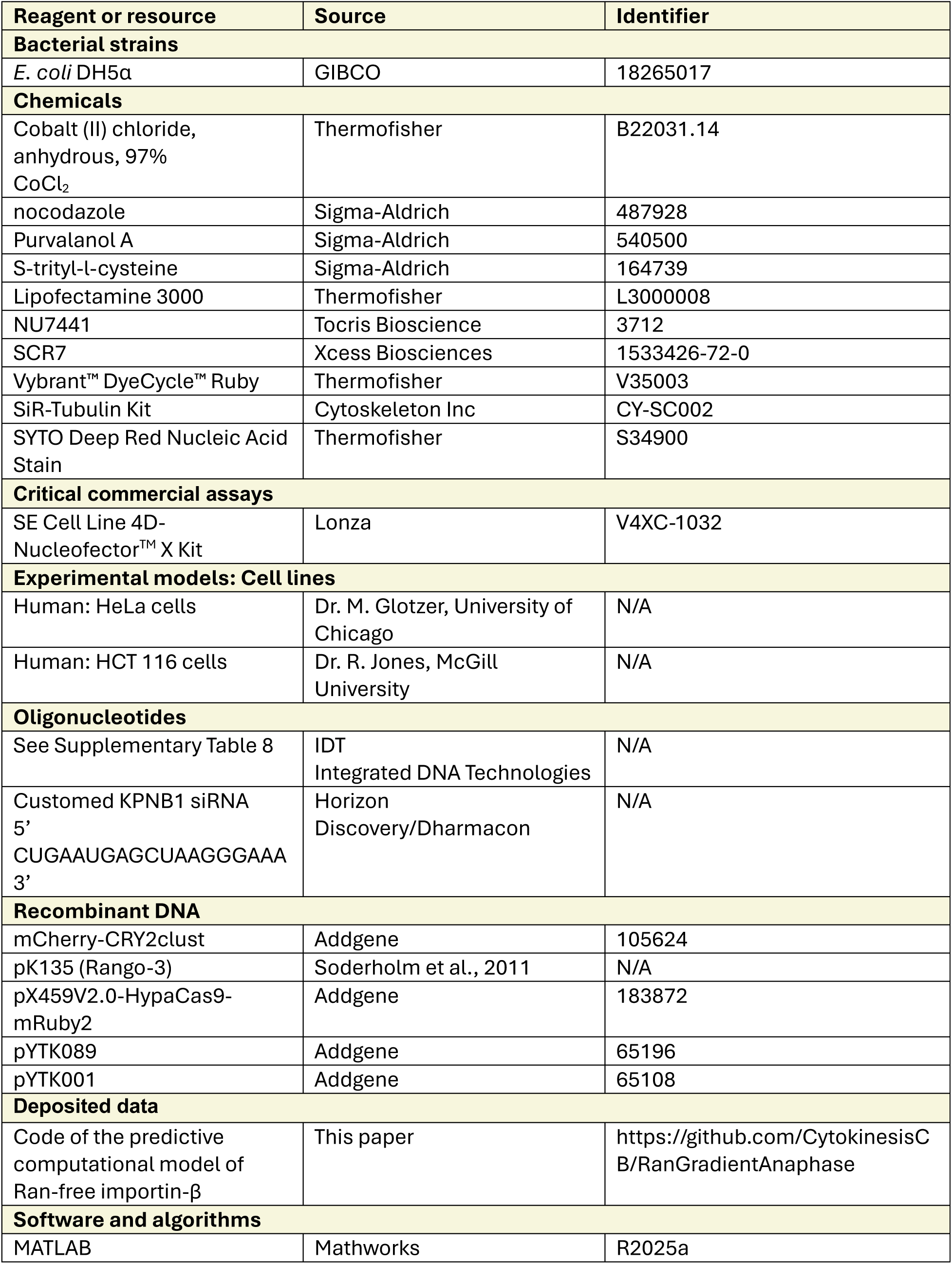
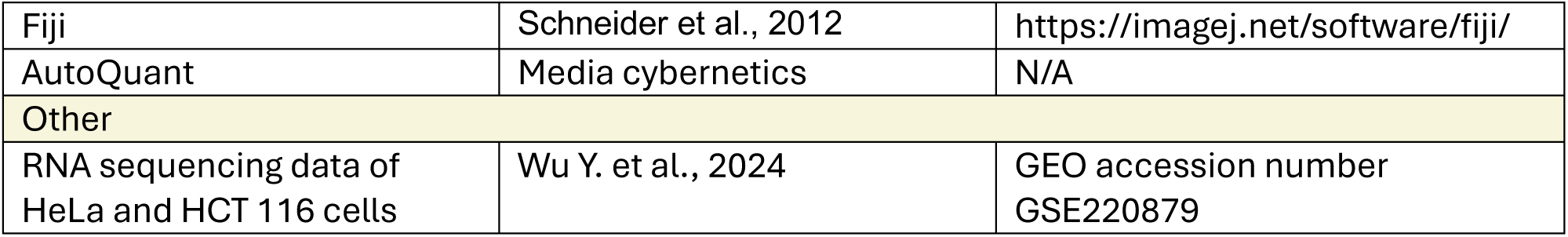

### Cell culture, transfection and drug treatment

HeLa cells were cultured in Dulbecco’s modified Eagle medium (DMEM; Wisent) media supplemented with 10% Cosmic calf serum (CCS; Cytoviva). HCT116 (p53-/-) cells were cultured in McCoy’s media (Wisent) supplemented with 10% CCS. The cells were maintained in 10 cm dishes in incubators at 37°C with 5% CO_2_. For plating, cells were counted using Countess 3 Automated Cell Counter (Thermofisher).

For transfection, HeLa cells were plated in DMEM media without antibiotics (penicillin and streptomycin: PS; Thermofisher) and transfected with DNA (cDNA and shRNA) or DNA and siRNA using Lipofectamine 3000 (Invitrogen) according to the manufacturer’s protocol. We modified the amount of DNA or siRNA to maintain molar ratios as recommended. HCT116 cells were transfected in suspension before plating. Cells were analysed 48h after transfection.

To induce monopolar cytokinesis, 2 μM of S-trityl-L-cysteine (STC; Millipore-Sigma) was added to cells for 8-16h, arresting them in prometaphase, after which Purvalanol A (Millipore-Sigma) was added to afinal concentration of 22.5 μM to inhibit Cdk1 for mitotic exit^5,21^. Cells were imaged immediately after Cdk1 inhibition. To arrest mitotic cells lacking microtubules, 100 nM of nocodazole (Sigma-Aldrich) was added to cells for 3–4h, then Purvalanol A was added to induce mitotic exit as described above.

To increase ploidy, HCT116 (p53-/-) cells were treated with 100-150 µM CoCl_2_ (Millipore-Sigma) for 48h as described previously^33^. The media was changed 3-4h before imaging. The FACSMelody cell sorter (BD Biosciences) was used to measure changes in ploidy in the population. After CoCl_2_ treatment, cells were dissociated and resuspended in Hank balanced salt solution (HBSS). They were then treated with 5 µM Vybrant™ DyeCycle™ Ruby (Invitrogen) for 30 minutes at 37°C with 5% CO_2_ to stain chromatin according to manufacturer’s instructions. The proportion of cells with different intensities were measured using the FACSMelody and the analysis was done using FlowJo (BD).

The cytosolic volume of HeLa cells was increased using hypotonic media. Cells were plated as described above, then the media was changed to DMEM diluted with 25% of sterile water and 10% of CCS or FBS. Cells were imaged after 12h of incubation.

### Cloning

mScarlet:anillin and rescue plasmids, pGG-mScarlet:Anillin-shANLN1 and pGG-mScarlet:Anillin-NLS-mutant-shANLN1, were generated using Golden Gate assembly^39^. Briefly, coding sequences of anillin and anillin with mutations in the C terminus NLS (887KK888-DE) were amplified from previously generated constructs^5,15^ and cloned into the pYTK001 backbone^40^ (Addgene #65108) by Golden Gate using BsmBI (New England Biolab), then they were assembled with mScarlet into a modified pX459V2.0-HypaCas9 vector^41^ (Addgene #108294) using BsaI (New England Biolab) for mammalian expression. A shRNA sequence targeting endogenous anillin, designed using previously described siRNAs, was also cloned into the modified pX459V2.0-HypaCas9 vector^5^.

mCherry-CRY2clust cDNA was obtained by PCR from mCherry-CRY2clust^24^ (Addgene plasmid #105624). Importin-β1 cDNA was amplified by PCR from a previously generated construct^5^. Three silent mutations (spanning E760-761L) were generated to make the cDNA sequence resistant to KPNB1 siRNA using PCR-based site-directed mutagenesis. Importin-β1 and mCherry-CRY2clust sequences were assembled using Golden Gate assembly as described above. All constructs were transformed into *E. coli* DH5α and sequenced using Plasmidsaurus. Primers are listed in Supplementary Table S8.

### Genome editing

The sgRNA spacer sequence used to endogenously tag KPNB1 was previously described by OpenCell CZ BIOHUB^42^. The sgRNA spacer was cloned into the pX459V2.0-HypaCas9-mRuby2 plasmid (Addgene plasmid #183872) as described previously^2,17^. To build the repair template, the homology arms were amplified from genomic DNA extracted from HeLa cells. mNeonGreen was amplified independently using primers that match part of the homology arms (15–51 bp). The repair template was assembled into the pYTK089 (Addgene plasmid #65196) vector using NEBuilder® HiFi DNA Assembly (New England Biolab). pX459-HypaCas9-mRuby2 and the repair template plasmid were transfected into HeLa cells using Lipofectamine 3000 as described above. Transfection in HCT 116 (p53-/-) was done by electroporation using the SE Cell Line 4D-Nucleofector^TM^ X Kit (Lonza) as per the manufacturer’s protocol. Cells were treated with NHEJ inhibitors NU7441 (2 µM; Tocris Bioscience) and SCR7 (1 µM; Xcess Biosciences) 4h before transfection and for 48h after transfection to improve editing eMiciency. Cells were monitored for the appearance of mNeonGreen fluorescence after ∼5 days.

Single cells were collected by sorting 7 days after transfection using the FACSMelody cell sorter (BD Biosciences). To do this, cells were dissociated and resuspended in FACS buMer composed of PBS with 1 mM EDTA, 25 mM HEPES pH 7.0 and 1% FBS. The gates were set to capture single fluorescent cells which were recovered in a 96-well plate containing DMEM or McCoy’s with 20% of CCS and penicillin-streptomycin (50 units ml−1 penicillin and 50 µg ml−1 streptomycin; Wisent). Cells were left to recover and were supplemented with media as needed.

Editing was confirmed by PCR and heterozygous clones were selected. PCR products were sequenced by Plasmidsaurus to confirm no alteration of KPNB1 and mNG sequence.

### Microscopy

For live-cell imaging, cells were seeded onto acid-etched round coverslips (25 mm, no. 1.5) in 6-well plates and incubated at 37°C with 5% CO_2_ for 24-72h. To visualize DNA, SYTO Deep Red Nucleic Acid Stain (Thermofisher) was added to afinal concentration of 0.25-2.5 μM. To visualize microtubules, SiR-tubulin (Cytoskeleton Inc) was added to afinal concentration of 1.25 pM. The coverslips were transferred to a magnetic chamber (Quorum) with 1 ml of media before imaging. Imaging was performed using an inverted Nikon Eclipse Ti microscope (Nikon) equipped with a Livescan Sweptfield scanner (Nikon), Piezo Z stage (Prior), IXON 879 EMCCD camera (Andor), and 488, 561 and 640 nm lasers (100mW, Agilent) using the 100×/1.45 NA objective. The cells were kept at 37°C and 5% CO_2_ during imaging in an INU-TiZ-F1 chamber (MadCityLabs). Z-stacks of 1 µm thickness were acquired every 1-2 min using NIS Elements software (Nikon, v.4.0). We also imaged cells using a Zeiss Axio Observer microscope (Zeiss) equipped with a Cicero spinning disk (CrestOptics), a Hamamatsu Flash 4.0 LT camera (Hamamatsu) and 488, 561 and 640 nm LDI-5 laser lasers (89North) using the 63×/1.4 NA oil immersion objective (Zeiss; pixel size 0.26μm). The cells were kept at 37°C and 5% CO_2_ during imaging. Z-stacks of 1 µm thickness were acquired every 1-2 min using Volocity software (Quorum). Images of cells stained with SiR-tubulin were acquired using a Nikon Super-Resolution Spinning Disk (SR-SD) comprised of an inverted Nikon Eclipse Ti2 microscope with a CSU-X1 spinning disk (Yokogawa), GATACA Live-SR unit, 60×/1.4 CFI PLAN APO VC oil immersion objective (Nikon), piezo Z stage (MadCityLabs), and Zyla camera (Andor). All image files were exported as TIFFs and opened with Fiji (NIH).

### Optogenetics

HeLa cells with endogenous mNeonGreen-tagged importin-β1 were seeded at 30% confluency onto acid-etched round coverslips (25 mm, no. 1.5) in 6-well plates. The next day, they were co-transfected with KPNB1 RNAis (5’ CUGAAUGAGCUAAGGGAAA 3’; Horizon), KPNB1:mCherry:CRY2 using Lipofectamine 3000 as described above. Cells were filmed 45-50h after transfection. CRY2 was activated at anaphase onset with 800ms of exposure to 15% 488nm laser, which also confirmed endogenous importin-β1 depletion. Z-stacks of 16 slices of 1 µm per slice were acquired over time. Cells with >70% depletion of endogenous importin-β1 were selected for analysis.

### Fluorescence Lifetime Microscopy (FLIM)

Cells seeded in 35 mm glass bottom dishes with 20 mm μ-wells (Cellvis) were transfected with pK135, which encodes Rango-3^28^. 12 hours after transfection, FLIM-FRET images were acquired using a Leica Stellaris-8 inverted laser scanning confocal microscope equipped with a HC PL APO CS2 100x/1.40 oil objective, and 80 MHz pulsed white light laser, a time-correlated HyD S detector, with LAS X software and a Falcon module for phasor analysis of photon arrival time. Cells were maintained at 37°C and 5% CO_2_ using a Tokai Hit stagetop incubator. Images of 512 x 512 pixels with a lateral size of 0.076 x 0.076 μm were acquired by unidirectional scanning with the white light laser set to 470 nm and 2% total intensity, an emission window of 490-600 nm and the detector set to photon counting mode, with a scan rate of 400 Hz, a pixel dwell time of 2.825 μs and 8 - 16x line accumulation, with transmitted light simultaneously acquired using a TransPMT. Photon arrival times for each pixel were calculated by the phasor plot, and false colouring for lifetime information was generated by drawing a line through the main pixel population in the phasor plot, using a blue-to-red colour scheme reflecting shorter to longer lifetimes (Supp. Fig. S3). Cells with similar expression levels were used for analysis as fluorescence intensity is proportional to Rango-3 concentration (Supp. Fig. S4 and Supp. Table S1).

### Image analysis and quantification

All images were opened in Fiji (v.2.3, NIH). Images of importin-β1 acquired by the Nikon Eclipse Ti microscope (Nikon) equipped with a Livescan Sweptfield scanner (Nikon) were deconvolved using AutoQuant (Media cybernetics). To reveal anillin’s cortical localization, linescans of anillin fluorescence intensity were performed at the desired time point on a max z-projection of three z-slices (1 μm per slice) by drawing a 3-pixel-wide line around the cortex of the cell. To calculate the breadth to size ratio, the breadth of the peak was measured as the number of pixels above 50% of the maximum value, which was subsequently divided by the length of the cell.

To determine the ratio of cortical to cytosolic anillin fluorescence, the background was first subtracted from the image, then three regions of interest (ROI) were drawn in the cytosol or at the cortex and averaged, then the cortical average value was divided by the cytosolic average value. To reveal subcortical importin-β1 localization, linescans of importin-β1 fluorescence intensity were performed at the desired time point on a sum z-projection of three z-slices (1 µm per slice) by drawing a 10-pixel-wide line around the cortex of the cell. Importin-β1 linescans were plotted together by normalizing them by their median. The breadth of importin-β1 was determined as per anillin but measuring the number of pixels above 60% of the peak maximum value.

To measure the correlation between enriched anillin and importin-β1, maximum z-projections of three z-slices (1 µm per slice) were used for anillin and sum z-projections of three z-slices (1µm per slice) were used for importin-β1. A mask was created and applied to each projection to define the zones of enrichment. The angle between the equatorial axis and the boundary of the breadth of enrichment was measured and plotted for each protein.

Measurements including the length and width of chromatin and the length and width of cells, spindle length and the distance between the segregating chromosomes, were done on images acquired using the Nikon Super-Resolution Spinning Disk (SR-SD). Spindle length was measured from centrosome to centrosome, and the distance between chromosomes was measured from the middle of one set of chromosomes to the middle of the other set. The means of each parameter from 10 cells were used for the computational model.

All data was exported as Excel (Microsoft)files for analysis. All graphs and statistics were generated using Prism (v.9.3, GraphPad). Statistical significance was tested using a Welch t-test.

### Ran-gradient modeling

The Ran-gradient model is based on a 1-D reaction-diffusion model from Caudron et al.^20^ converted into a model with 2-D geometry. The partial differential equations were solved using the PDE toolbox of MATLAB (R2025a).

#### Description of the reaction and diffusion coefficients

The cell was divided into two zones: cytosol and chromatin (Supp. Fig. S5 and S6). We assume that the guanine nucleotide exchange from Ran-GDP to Ran-GTP happens only in the chromatin zone since the guanine nucleotide exchange factor (GEF), RCC1, is bound to histone and does not diffuse (Moore et al., 2002). We assume that Ran-GTP, Ran-GDP, GTP and GDP can freely diffuse through all zones. The hydrolysis of Ran-GTP into Ran-GDP and the interaction between Ran-GTP and importin-β take place in the cytosol. The RanBP1 interaction with Ran-GTP can be neglected due to the rapid hydrolysis of GTP, according to Caudron et al.^20^. We used the same diffusion coeMicients as well as reaction rates and kinetic constants as Caudron et al.^20^ (Supp. Table S2 and S3).

#### Description of the equations

The reactions occur in a 2-D space representing a cell at metaphase or anaphase. The concentration of proteins depends on their position (x and y) within the cell and time. The reaction-diffusion equation for a concentration u (x,y,t) is defined as the following:

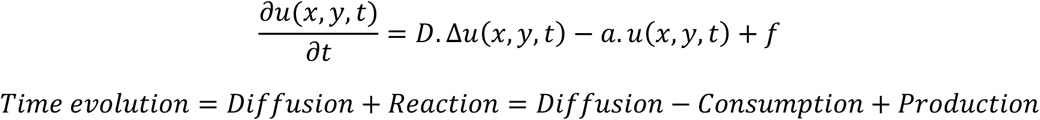

#### In the chromatin zone

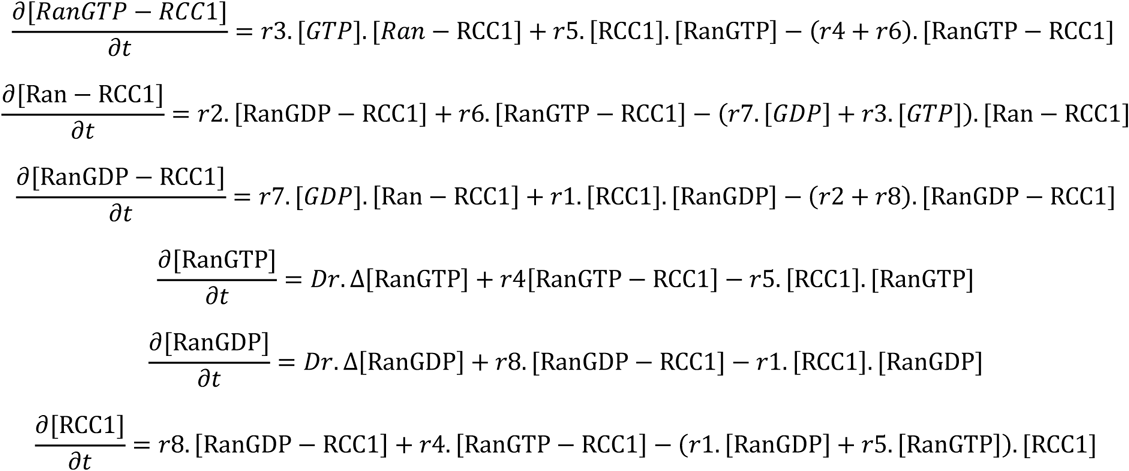

#### In the cytosol zone

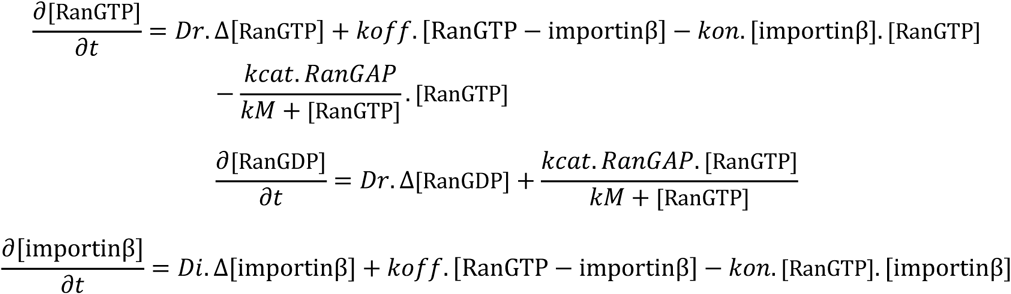

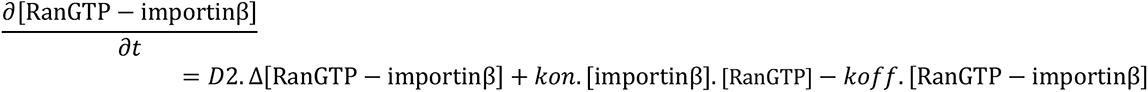

The calculations were done using the Partial Differential Equation (PDE) solver of Matlab®.

#### Description of geometry and boundary conditions

There is a reflective condition (no flux) at the edge of the cell or chromatin for proteins that cannot escape their respective zone (Supp. Table S2).

The parameters used for the equations above come from measurements of metaphase and anaphase HeLa and HCT116 cells (Supp. Figure S6). These include cell and chromatin length and width, as well as spindle length. Our measurements revealed that the cell and chromatin volume did not vary significantly between metaphase and anaphase cells, therefore we assumed no change in concentration between metaphase and anaphase. Volume errors were calculated using standard analytical propagation: 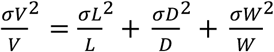

The mesh is shown in Supplementary Figure S6. The spatial steps were chosen to be small enough to capture the important features of the solution, but not too small that the computation becomes excessive and difficult to run on a standard computer. The maximum size of the mesh in the cytosol is 1 µm, while in the chromatin zone, it is 0.1 µm. The mesh was defined to include 15 grid points across the chromatin width to prevent numerical instability due to boundary conditions or the creation of a sharp gradient. 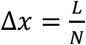, where L is the smallest distance used, and N is the number of grid points.

#### RNA sequencing analysis and estimation of the initial conditions of the computational model

The total cellular concentrations of Ran, RanGAP and importin-β in HeLa cells are 6 μM, 0.4 μM and 3 μM, respectively, which were calculated in previous studies^43,44^. Assuming that RanGAP and importin-β1 are restricted to the cytosol, their local concentrations are 0.7 μM and 5.25 μM, respectively. We assumed that RCC1 concentration is 0.24 μM^20^. However, since RCC1 is localized to chromatin, it is ∼10 to 100-fold more concentrated at this location compared to the cytosol^43,44^.

Since such estimates of protein concentrations are not available for HCT 116 cells, we re-analysed published RNA sequencing data for HeLa and HCT 116 cells ^31^ with GEO2R (GEO accession number: GSE220879) to measure the expression of RAN, RCC1, RANGAP1 and KPNB1. We used their relative expression between HCT 116 and HeLa cells to approximate protein concentrations in HCT 116 cells (Supp. Figure S7). We acknowledge that RNA sequencing represents only the levels of mRNA and there could be differences in protein levels. However, mRNA can be used as a proxy for protein. First, we compared the fold-change to Ran in HeLa cells.

Assuming that [*Ran*]*_total_* = 6 *μM* in HeLa cells:

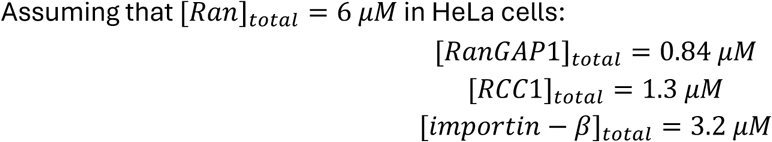

The total concentrations are similar between HeLa and HCT 116 cells with some small but significant changes in RanGAP1 and KPNB1 as indicated in (Supp. Figure S7). Using these values and the measurements of cell volume (Supp. Table S4), we estimated the concentrations of Ran, RanGAP1, RCC1 and importin-β in HCT 116 cells (Supp. Table S5). Note that errors of concentrations were calculated using standard analytical propagation: 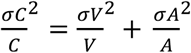

Deterministic computational models are imperfect as their results are based on input. We are forced to make assumptions due to technical limitations, but these assumptions must be physically plausible. The calculated errors for the initial conditions provide a range for each parameter to be tested experimentally (Supp. Figure S8). In addition, we evaluated the impact of each parameter on the system, which helped us to make the adjustments shown in Supp. Figure S9.

#### Estimation of equilibrium

The time steps needed to be small enough to capture the dynamics of the system, but not so small that they require unnecessary computational needs. For 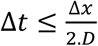, D is the greatest diffusion rate. The dynamics of the system can be resolved using a time step of 0.000227 seconds.

The model starts from a state of non-equilibrium: all of Ran is inactive and free, RCC1 and RanGAP1 are free within their respective zones, and all importin-β is free. The model is stopped when the system has reached an equilibrium. We picked t_end_ as the difference between t_end_ and t_end-1_ is nul.

#### Establishing changes in parameters that impact the cytosol to chromatin ratio

To model HeLa cells with a decrease in chromatin-to-cell-size ratio based on an increase in cytosol volume, we adjusted the length and width of the cell (Supp. Table S6). We did not change the distance between chromatin or chromatin shape. The concentrations of proteins were adjusted accordingly (Supp. Table S7).

To model HCT116 cells with an increase in chromatin-to-cell-size ratio based on an increase in ploidy, we doubled the DNA content and increased both the cell and chromatin volumes by 1.5 (Supp. Table S6). The concentrations of proteins were adjusted accordingly (Supp. Table S7).

## Resource availability

The MATLAB code for the predictive computational model of Ran-free importin-β is available at: https://github.com/CytokinesisCB/RanGradientAnaphase

## Acknowledgments

We thank M. Husser and I. Ozugergin for their help with the gene editing method. We thank C. Law for help with imaging studies as part of the Centre for Microscopy and Cellular Imaging at Concordia University. We thank J. Ryan for help with FLIM-FRET images collecting and processing at the Advanced BioImaging Facility (ABIF) at McGill University (RRID: SCR_017697). This work was funded by NSERC.

## Author contributions

K.L. conceptualized, designed and conducted the experiments for Figure 6D, E and G. Perlman conducted experiments for Figure 6D, E and G. I.T. performed experiments for Figure 3A and 6A.

C.R.B. wrote the manuscript including all figures, conceptualized, designed and conducted experiments including analysis and entirely did the computational modelling. A.P. supervised, edited the manuscript and conceptualized this study.

## Supplemental information

Document S1. Supplementary Figures 1–11 and Supplementary Tables 1-7. Supplementary Table 8 – Primers’ list

