## Supplemental Files for "Importin-β1 functions as a chromatin sensor to position the contractile ring for cytokinesis"

**Supplementary Information**


- Supplementary Figure S1. Genotyping of importin-β1:mNG tagged cells
- Supplemental Figure S2. Anillin localizes differently in mitotic HeLa and HCT 116 cells
- Supplementary Figure S3. Method used to image and evaluate the fluorescence lifetime
- Supplementary Figure S4. Fluorescence lifetime is dependent on the levels of Rango-3
- Supplementary Figure S5. Reactions used to calculate the Ran gradient in the predictive model.
- Supplementary Figure S6. Cell measurements used in the predictive model
- Supplementary Figure S7. RNA-seq data analysis for Ran and Ran regulators in HeLa and HCT 116 cells
- Supplementary Figure S8. Method for model refinement
- Supplementary Figure S9. Impact of changing the levels of Ran or Ran regulators on the predicted Ran-free importin-β1 concentration
- Supplemental Figure S10. Generation of hypotonic HeLa cells and hyperploid HCT 116 cells
- Supplementary Figure S11. The spindle scales with cell size
- Supplementary Table S1. Rango-3 fluorescence intensity in individual HeLa cells
- Supplementary Table S2. Diffusion coefficients for Ran and Ran regulators
- Supplementary Table S3. Reaction rates and kinetic constants for Ran and Ran regulators
- Supplementary Table S4. Measurements of components in metaphase and anaphase HeLa and HCT 116 cells
- Supplementary Table S5. Concentrations of Ran and Ran regulators in HeLa and HCT 116 cells
- Supplementary Table S6. Initial conditions for the predictive Ran-free importin-β1 gradient model
- Supplementary Table S7. Measurements for anaphase hypotonic HeLa and hyperploid HCT116 cells
- Supplementary Table S8. List of primers used for cloning

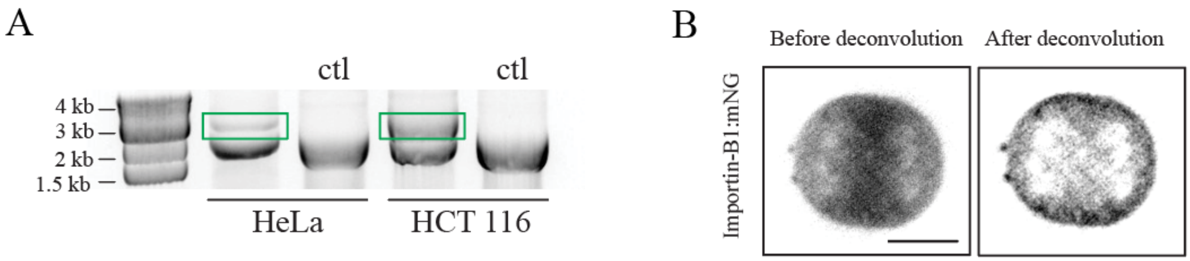


***Supplementary Figure S1. Genotyping of importin-β1:mNG tagged cells.*** *(A) An image shows an agarose gel with PCR-amplified DNA for the KPNB1 locus and mNG insertion in HeLa and HCT 116 cells. Amplification of clones with the mNG insertion produced a 3kb fragment (green box), while non-tagged control cells produced a 2kb fragment. Note that the intensity of the mNG-inserted fragment in HeLa cells indicates that the clone is heterozygous, while the HCT 116 clone is homozygous. (B) Images show importin-β1:mNG in an anaphase HeLa cell before (left) and after deconvolution (right). The scale bar is 10 µm.*

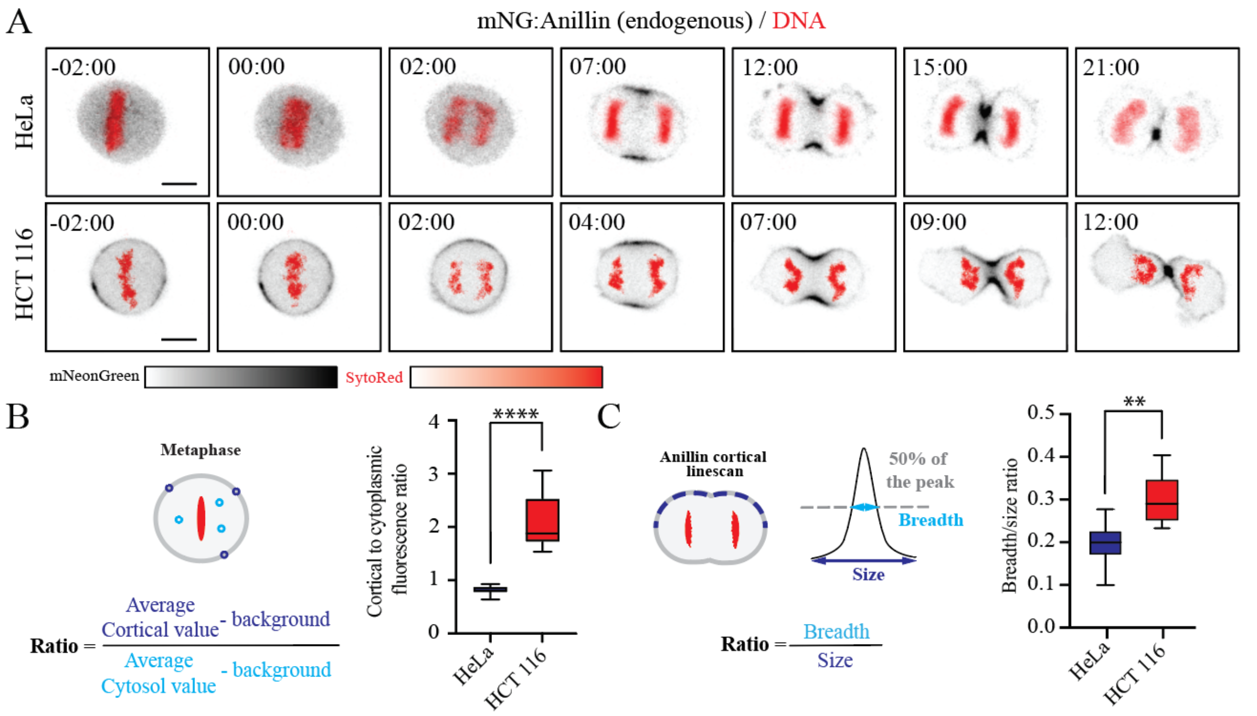


***Supplemental Figure S2. Anillin localizes differently in mitotic HeLa and HCT 116 cells.*** *(A) Timelapse images show endogenous anillin tagged with mNG (greyscale) co-stained for chromatin (Hoechst; red) in HeLa (top) and HCT 116 cells (bottom). The colour bar indicates the intensity of signal from light (low levels) to bright (high levels). Time is in minutes, and the scale bars are 10 µm. (B) A cartoon cell shows how the ratio of cortical to cytosolic anillin fluorescence was measured. Three regions of interest (ROI) were drawn in the cytosol or at the cortex and averaged, then the cortical average value was divided by the cytosolic average value. A bar graph shows the average cortical vs. cytoplasmic fluorescence ratio of anillin in metaphase HeLa (n=17) and HCT 116 cells (n=20; ****: $p \leq 0.0001$). (C) A schematic shows how the breadth of anillin in anaphase cells was measured using linescans, and the number of pixels >50% of the maximum peak value. The breadth was then divided by the length of the line to calculate the ratio of breadth to length (size). The graph shows the average ratio of breadth to size of anillin (>50% peak signal) in HeLa (n=17) and HCT 116 cells (n=20; **: $p \leq 0.005$).*

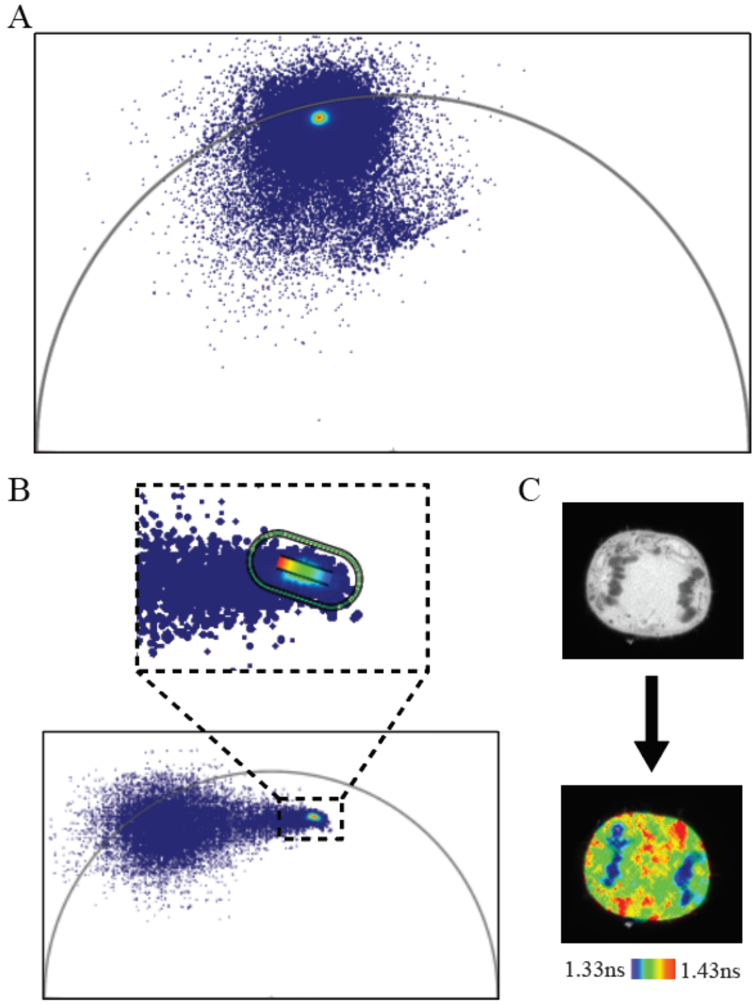


***Supplementary Figure S3. Methods used to image and evaluate fluorescence lifetime.*** *(A) A graph shows phasor plots of EGFP, which represents the fluorescence decay for each pixel. Mono-exponential decay occurs along the semi-circle. Signals on the left indicated higher lifetime compared to those on the right. Multi-exponential decay occurs inside the semi-circle. Signals closer to the centre are higher in complexity and reflect a mix of fluorophores. The density cloud indicates where the signal is located in the semi-circle. (B) A graph shows the phasor plot of a HeLa cell with Rango-3 at anaphase. The picture on top is a zoomed-in image of the signal. A mask was created (coloured line) on the image. (C) Images show Rango-3 fluorescence in the cell (top) and the measured lifetimes (bottom; shorter; blue to longer; red).*

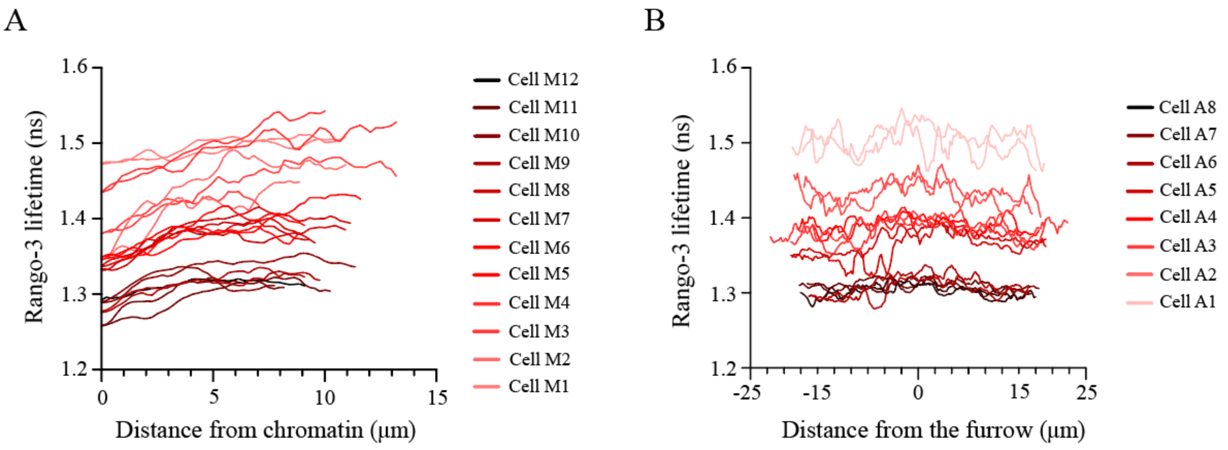


***Supplementary Figure S4. Fluorescence lifetime is dependent on the levels of Rango-3****. (A) A graph shows Rango-3 lifetime measurements in nanoseconds along a line drawn from chromatin to the cortex in metaphase HeLa cells (n=12). (B) A graph shows Rango-3 lifetime measurements in nanoseconds along the cortex of anaphase HeLa cells from pole to pole (n=8). The color scale is based on the intensity of Rango-3 fluorescence from dark (high levels) to light (low levels).*

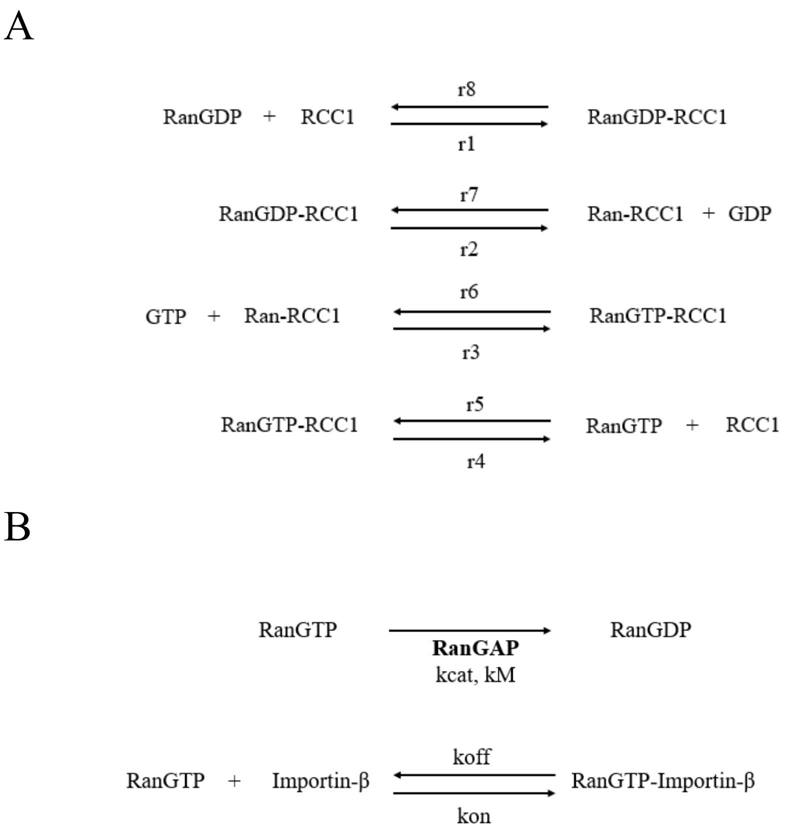


***Supplementary Figure S5. Reactions used to calculate the Ran gradient in the predictive model.*** *(A) Reactions for the RCC1-mediated guanine nucleotide exchange between Ran-GTP and Ran-GDP. These equations were applied to the chromatin zone. The reaction rates and kinetic constants are listed in Table S2. (B) Reactions for the hydrolysis of Ran-GTP by RanGAP, and Ran-GTP interaction with importin-β. These equations were applied to the cytosol zone.*

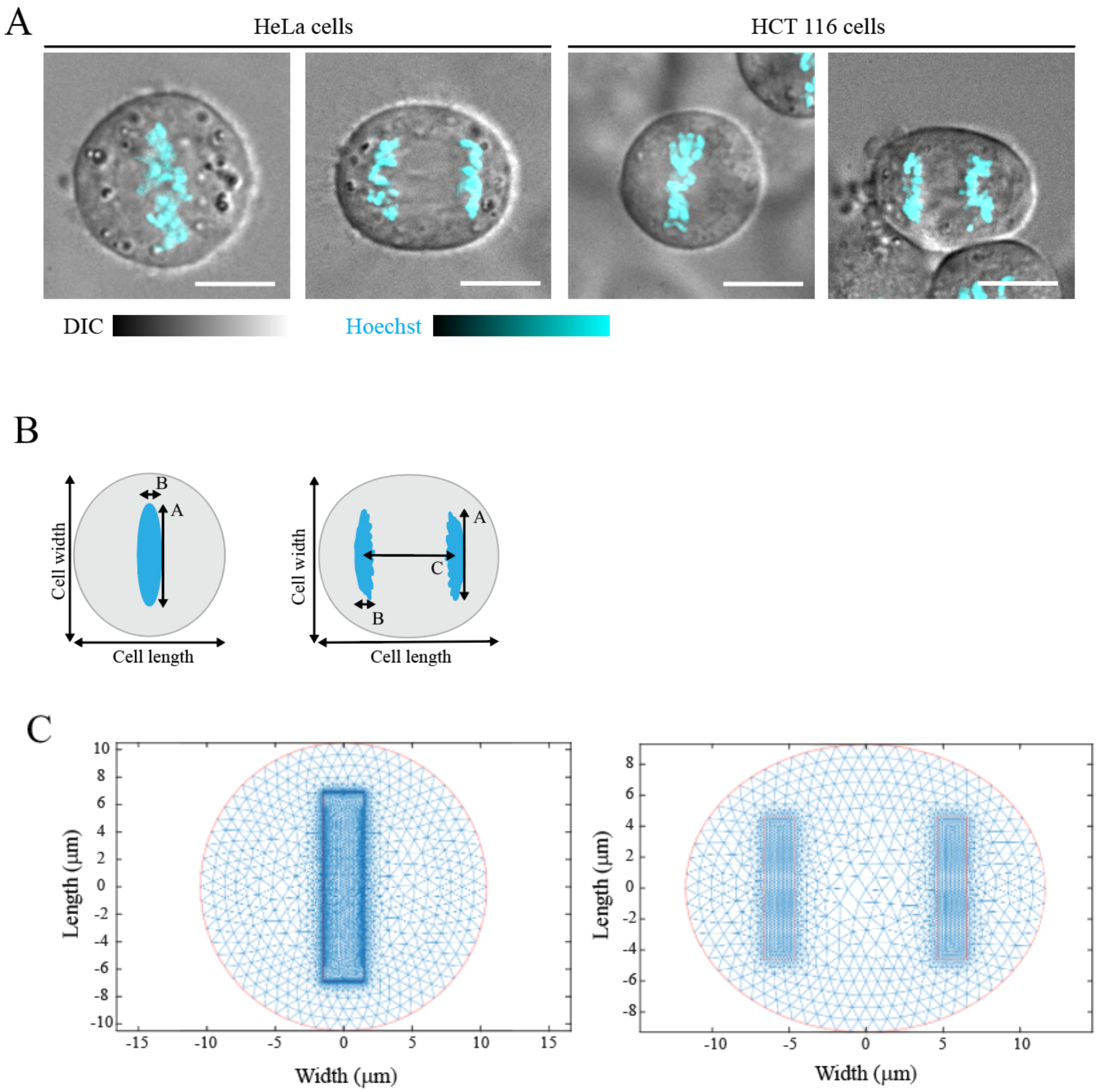


***Supplementary Figure S6. Cell measurements and mesh regions used in the predictive model.*** *(A) Images show DIC metaphase and anaphase HeLa and HCT 116 cells stained with SiR-Tubulin (red) and Hoechst (chromatin, cyan). The colour scales indicate fluorescence intensity from dark (low levels) to bright (high levels). The scale bars are 10 μm. (B) A schematic shows how cellular components were measured (A indicates chromatin length; B indicates chromatin width; C indicates the distance between sister chromatids). (C) The mesh used for the 2D model is shown for a HeLa cell at metaphase (left; n=10) and anaphase (right; n=10). The red line shows the outer boundary of the cell, while the blue hatch marks indicate the equations solved in the cytosol, and the lines show those near chromatin.*

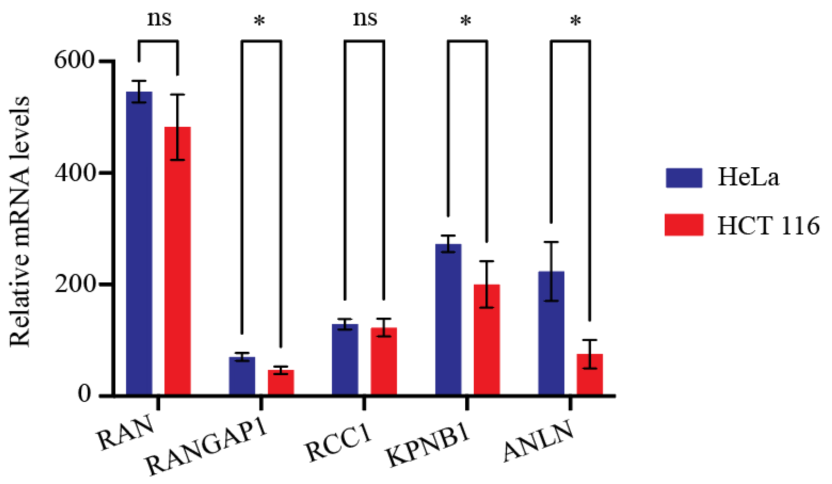


***Supplementary Figure S7. RNA-seq data analysis for Ran and Ran regulators in HeLa and HCT 116 cells.*** *The graph shows normalized mRNA expression levels for Ran, RanGAP1, RCC1, KPNB1 and ANLN in HeLa (blue) and HCT 116 cells (red; * $p < 0.05$; ns = non-significance) using the RNA sequencing data from Wu et al., 2024 (GSE220879).*

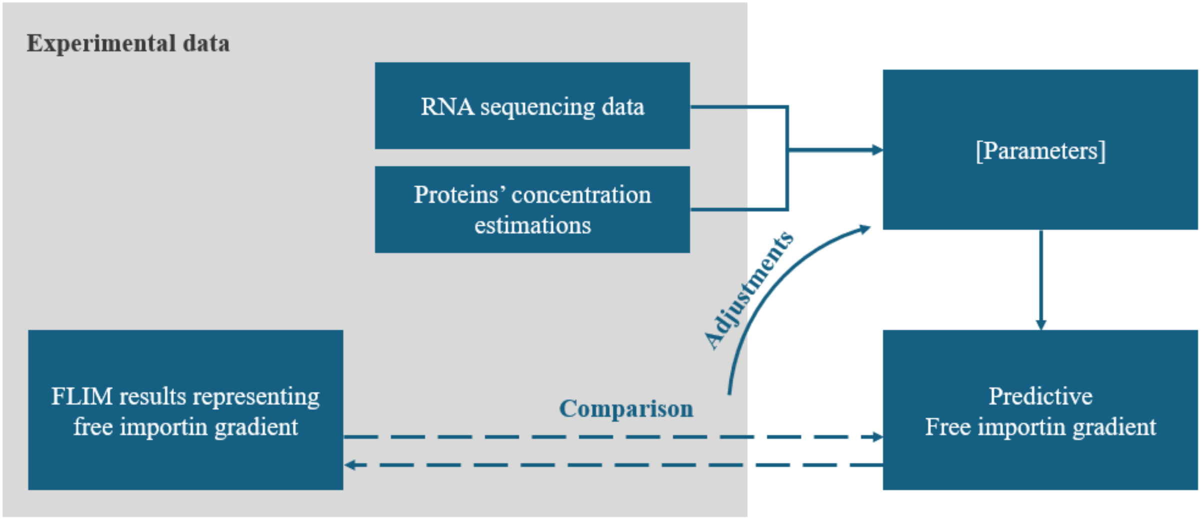


***Supplementary Figure S8. Method for model refinement.*** *A cartoon shows the method used to find conditions that match experimental findings. Parameters implemented in the predictive model include cell measurements, kinetic constants, reaction rates and protein concentrations. These values were taken from published data or measured and adjusted in the model using this workflow.*

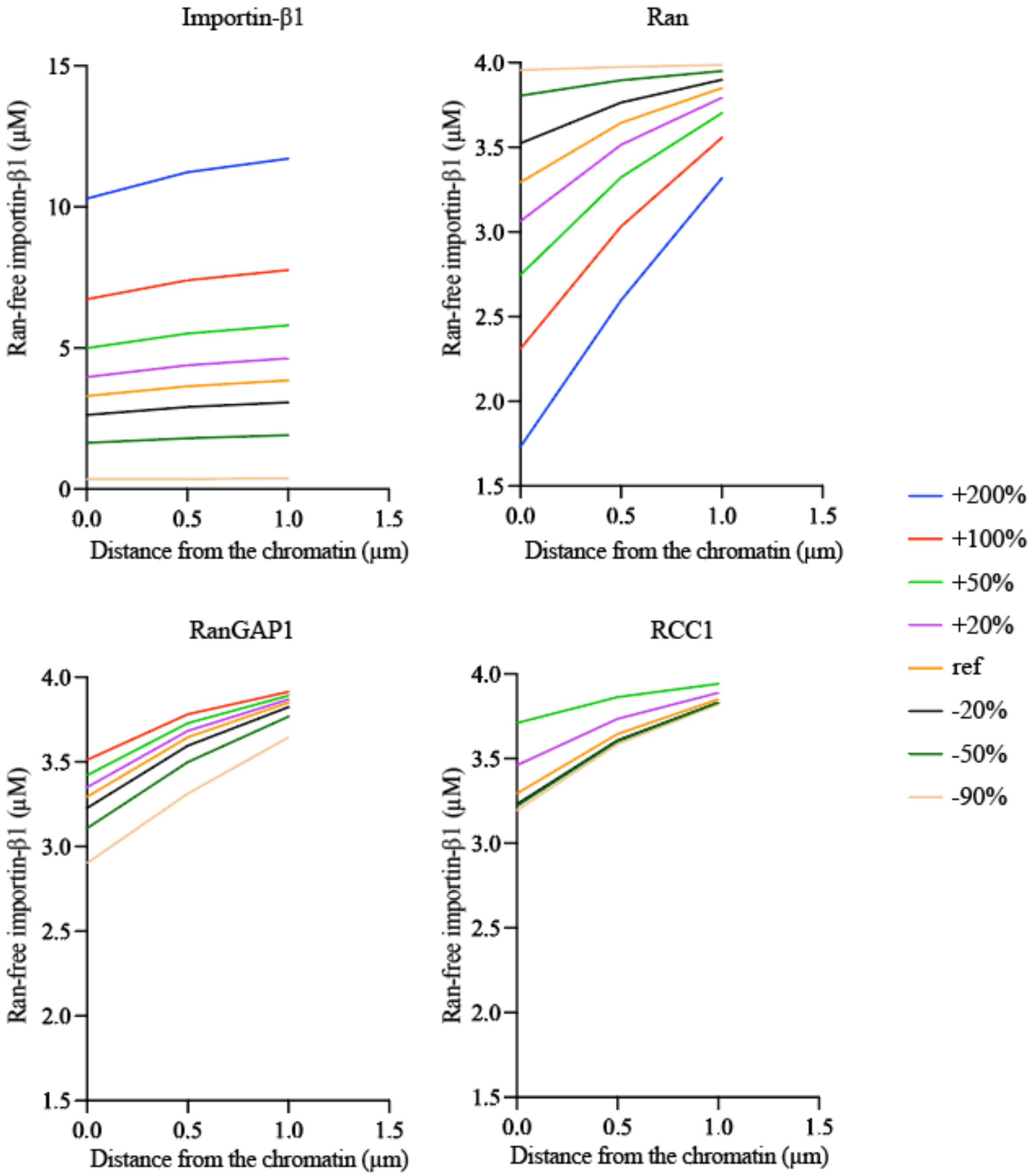


***Supplementary Figure S9. Changing the levels of Ran or importin-β1 had a greater impact on the predicted Ran-free importin-β1 concentration compared to RanGAP1 or RCC1.*** *Graphs show the Ran-free importin-β1 concentration (µM) in metaphase cells from chromatin to the cortex in response to changes in the levels of Ran, RCC1, RanGAP and importin-β1. The conditions vary between -90% and 200% in relation to the concentrations calculated based on the RNA sequencing data (ref; reference).*

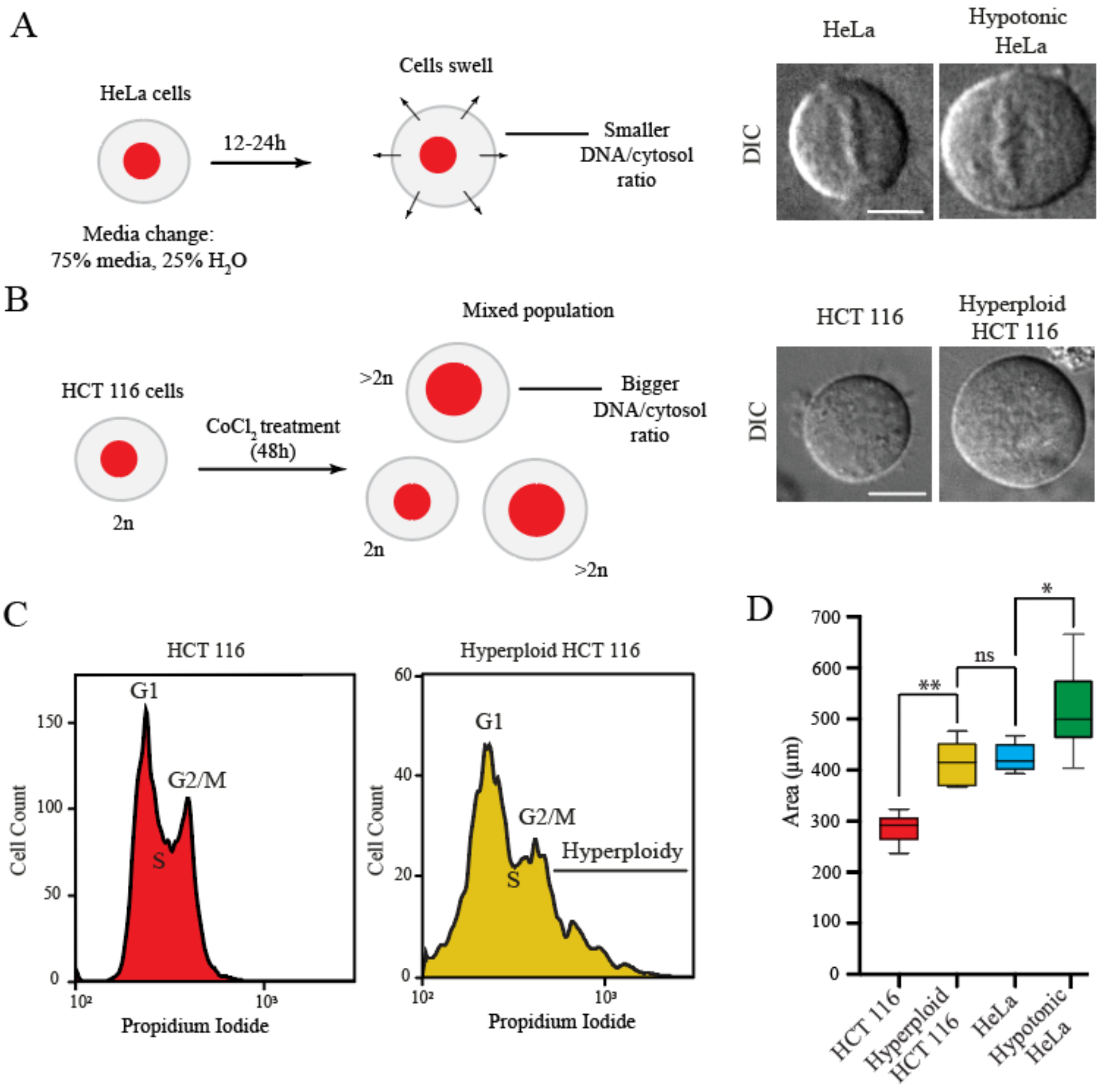


***Supplemental Figure S10. Generation of hypotonic HeLa cells and hyperploid HCT 116 cells.*** *(A) The schematic on the left shows the workflow to generate hypotonic HeLa cells. Cells were cultured in 75% hypotonic media supplemented with FBS for 12h resulting in large HeLa cells. The DIC images show a metaphase HeLa cell in isotonic media (left) and in hypotonic media (right). The scale bar is 10 µm. (B) The schematic on the left shows the workflow to generate hyperploid HCT 116 cells. Cells were treated with* $CoCl_2$ *for 48h resulting in a mixed population of diploid and hyperploid cells. The DIC images on the right show a metaphase diploid HCT 116 cell (left) and hyperploid cell (right). The scale bar is 10 µm. (C) The histograms show the DNA content of HCT 116 cells (left) and after treatment to increase ploidy (right). DNA was stained with propidium iodine. The cell cycle stages are indicated in the graphs. (D) The graph shows the area of metaphase HCT 116, hyperploid HCT 116, HeLa, and hypotonic HeLa cells (n=15 for each category;* $*p<0.005$*;* $**p<0.0001$*; n.s., no significance).*

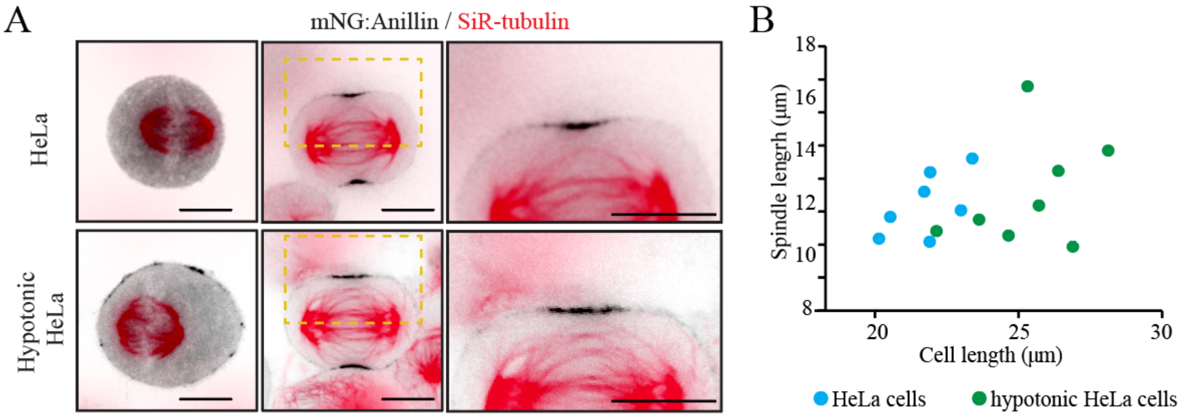


***Supplementary Figure S11. The mitotic spindle scales with cell size.*** *(A) Images show HeLa cells and hypotonic HeLa cells at metaphase (left) and anaphase (right) with endogenous anillin tagged with mNG (greyscale) and co-stained for tubulin (SiR-Tubulin; red). The images on the right are zoomed-in views of the yellow boxes from anaphase cells. The scale bars are 10 µm. (B) A graph shows the correlation between the spindle length and cell length in metaphase cells. The dots show individual cells (blue, HeLa cells, n=7; green, hypotonic HeLa cells, n=8).*

***Supplementary Table S1. Rango-3 fluorescence intensity in individual HeLa cells****. Values indicate the average fluorescence intensity of Rango-3 in metaphase and anaphase HeLa cells. Grey boxes indicate the cells used for analysis in the main text.*

| Metaphase cells | Fluorescence intensity | Anaphase cells | Fluorescence intensity |
|---|---|---|---|
| Cell M1 | 195.12 | Cell A1 | 247.006 |
| Cell M2 | 205.1 | Cell A2 | 501.532 |
| Cell M3 | 531.982 | Cell A3 | 626.57 |
| Cell M4 | 586.031 | Cell A4 | 1038.823 |
| Cell M5 | 556.447 | Cell A5 | 1283.053 |
| Cell M6 | 603.82 | Cell A6 | 1685.009 |
| Cell M7 | 1142.071 | Cell A7 | 2219.852 |
| Cell M8 | 1777.825 | Cell A8 | 2052 |
| Cell M9 | 2231.665 | | |
| Cell M10 | 2122.818 | | |
| Cell M11 | 1632.743 | | |
| Cell M12 | 4068.93 | | |

***Supplementary Table S2. Diffusion coefficients for Ran and Ran regulators****.*

| **Symbol in the code** | **Proteins** | **Localization** | **Diffusion coefficient name** | **D (μm2/sec) in the cytoplasm** |
|---|---|---|---|---|
| u1 | RanGTP-RCC1 | Chromatins, immobile | NA | NA |
| u2 | Ran-RCC1 | Chromatins, immobile | NA | NA |
| u3 | RanGDP-RCC1 | Chromatins, immobile | NA | NA |
| u4 | RanGTP | Chromatins/cytosol/spindle | Dr | 22 |
| u5 | RanGDP | Chromatins/cytosol/spindle | Dr | 22 |
| u6 | RCC1 | Chromatins, immobile | NA | NA |
| u7 | Importin-β | Cytosol/spindle | Di | 14 |
| u8 | RanGTP-ImportinB | Cytosol/spindle | D2 | 13 |

***Supplementary Table S3. Reaction rates and kinetic constants for Ran and Ran regulators.***

| **In chromatins** | **In cytosol** |
|---|---|
| r1 = 74.4 $\mu M^{-1}.s^{-1}$<br>r2 = 21.1 $s^{-1}$<br>r3 = 0.65 $\mu M^{-1}.s^{-1}$<br>r4 = 55 $s^{-1}$<br>r5 = 102 $\mu M^{-1}.s^{-1}$<br>r6 = 19 $s^{-1}$<br>r7 = 11.4 $\mu M^{-1}.s^{-1}$<br>r8 = 55 $s^{-1}$ | kcat = 10.6 $s^{-1}$<br>kM = 0.7 μM<br>kcat2 = 10.8 $s^{-1}$<br>kM2 = 0.1μM<br>kon = 0.45 $\mu M^{-1}.s^{-1}$<br>koff = 0.00045 $s^{-1}$ |

***Supplementary Table S4. Measurements of components in metaphase and anaphase HeLa and HCT 116 cells.*** *Measurements were made using images (n=10 cells each).*

| | HeLa cells | | HCT 116 cells | |
|---|---|---|---|---|
| | Metaphase | Anaphase | Metaphase | Anaphase |
| Distance between the chromatin (μm) | NA | 11.5±0.7 | NA | 8.4±0.5 |
| Length of chromatin (μm) | 13.8±1 | 9.5±0.7 | 12±1 | 7.5±0.6 |
| Width of chromatin (μm) | 3±0.5 | 2±0.3 | 3±0.5 | 2±0.1 |
| Depth of chromatins (μm) | 20±1.6 | 19±2.4 | 24±0.6 | 19±1.6 |
| Cell length (μm) | 21±3.9 | 22.6±1.1 | 17.4±2.4 | 18.8±1.19 |
| Cell width (μm) | 21±3.9 | 18.6±1.3 | 17.4±2.4 | 15.6±0.76 |
| Cell depth (μm) | 21±1.4 | 21±2.7 | 24±2.5 | 22±1.9 |
| Cell volume ($\mu m^3$) | 4920 ± 900 | 4850 ± 750 | 3920 ± 780 | 3440±320 |
| Chromatin volume ($\mu m^3$) | 830 ± 110 | 720 ± 140 | 670 ± 79 | 570 ± 82 |

***Supplementary Table S5. Concentrations of Ran and Ran regulators in HeLa and HCT 116 cells.*** *Concentrations are in μM and were evaluated using published RNA sequencing data.*

| | HeLa cells | HCT 116 cells |
|---|---|---|
| Ran (total concentration) | 6 ±1.6 | 6.6±2.4 |
| RanGAP1 (cytosol) | 2.5 ± 0.3 | 0.9±0.4 |
| RanGAP1 (cytosol) | 0.3±0.1 | 0.9 ± 0.4 |
| RCC1 (chromatins' zone) | 7.7±1.2 | 7±1.6 |
| Importin-β1 (cytosol) | 3.8±1.3 | 3.8±1.8 |

***Supplementary Table S6. Initial conditions for the predictive Ran-free importin-β1 gradient model.*** *Concentrations are in μM.*

| | HeLa cell | HeLa cell with increased cytosol | HCT 116 cell | Tetraploid HCT 116 cell |
|---|---|---|---|---|
| Ran (total) | 6.7 | 6.2 | 9.1 | 12.1 |
| RanGAP1 (cytosol) | 2.5 | 2.5 | 0.9 | 1.2 |
| RanGAP1 (cytosol) | 0.3 | 0.16 | 0.9 | 1.2 |
| RCC1 (chromatin zone) | 8.7 | 8.7 | 8.7 | 11.6 |
| Importin-β1 (cytosol) | 4 | 3.7 | 2.75 | 3.7 |

***Supplementary Table S7. Measurements for anaphase hypotonic HeLa and hyperploid HCT116 cells.***

| | Large HeLa cells | Hyperploid HCT116 cells |
|---|---|---|
| Distance between the chromatins (μm) | 11.5 | 9.6 |
| Length of chromatins (μm) | 9.5 | 8.6 |
| Width of chromatin (μm) | 2 | 2.3 |
| Cell length (μm) | 24 | 21.5 |
| Cell width (μm) | 19.1 | 17.9 |
| Cell depth (μm) | 22 | 25 |
| Total cell volume (μm$^3$) | 5380 | 5160 |
| Chromatins Volume (μm$^3$) | 830 | 860 |

***Supplementary Table S8. List of primers used for cloning.***

| **Final Plasmids** | **Gene target / name** | **Fwd/ Rev** | **Sequence** |
|---|---|---|---|
| KPNB1:mCherry:CRY2 | KPNB1 | FRW | CTGAATGAATTGAGGGAAAGCTGCTTGGAAGCCTATACT |
| | KPNB1 | REV | TTTCCCTCAATTCATTCAGATAATCCACCATGTCATAGTCTGACTT |
| | mCherry:CRY2clust | FRW | GCATCGTCTCATCGGTCTCAGGGGGTTCAGGAGGGTCTatggtgagcaagggcgaggagg |
| | mCherry:CRY2clust | REV | ATGCCGTCTCAGGTCTCActcatcagttatctagatccggtggatcc |
| | KPNB1 | FRW | GCATCGTCTCATCGGTCTCAccATGGAGCTGATCACCATTCTCGAGA |
| | KPNB1 | REV | ATGCCGTCTCAGGTCTCACCCCCTTCAGTTTCCTCAGTTCTTTTGT |
| p63R | N/A | FRW | CACCGAGGCAAGATCTTACCAAGCT |
| | N/A | REV | AAACAGCTTGGTAAGATCTTGCCTC |
| KPNB1 repair template | Genomic DNA | FRW | GACAGCTGCATCAGACCttctc |
| | Genomic DNA | REV | AGCTTGGTTCTTCAGTTTCCTCAG |
| | Genomic DNA | FRW | TGGTAAGATCTTGCCTCCACTGT |
| | Genomic DNA | REV | aggaggctgaggtgggtg |
| | mNeonGreen | FRW | AAAGAACTGAGGAAACTGAAGAACCAAGCTGGAGGTTCAGGAGGCTCCatggtgagcaagggcgag |
| | mNeonGreen | REV | AAAGAGGACAGTGGAGGCAAGATCTTACCActtgtacagctcgtccatgcc |
| | pYTK089 | FRW | ttttaggtgatccacccacctcagcctcctgattatcaaaaaggatcttcacctagatccttt |
| | pYTK090 | REV | aaaagagagagagaaGGTCTGATGCAGCTGTCacggttatccacagaatcagggg |
| pGG_mScarlet:Anillin, pGG-mScarlet:Anillin-shANLN1 and pGG-mScarlet:Anillin(NLSmutant)-shANLN1 | Anillin | FRW | GCATCGTCTCATCGGTCTCAGTCTatggatccgtttacggagaaactg |
| | Anillin | REV | ATGCCGTCTCAGGTCTCActcaaggctttccaataggtttgtagcaa |
| | N/A | FRW | caccgGGCGATGCCTCTTTGAATAAATTCAAGAGATTTATTCAAAGAGGCATCGCC |
| | N/A | REV | aaaaGGCGATGCCTCTTTGAATAAATCTCTTGAATTTATTCAAAGAGGCATCGCCc |
| | mScarlet | FRW | GCATCGTCTCATCGGTCTCAccatGGTGAGCAAGGGCGAG |
| | mScarlet | REV | ATGCCGTCTCAGGTCTCAAGACCCTCCTGAACCCCCCTTGTACAGCTCGTCCATGCC |
